# Glioma-Induced Alterations in Excitatory Neurons are Reversed by mTOR Inhibition

**DOI:** 10.1101/2024.01.10.575092

**Authors:** Alexander R. Goldberg, Athanassios Dovas, Daniela Torres, Sohani Das Sharma, Angeliki Mela, Edward M. Merricks, Markel Olabarria, Leila Abrishami Shokooh, Hanzhi T. Zhao, Corina Kotidis, Peter Calvaresi, Ashwin Viswanathan, Matei A. Banu, Aida Razavilar, Tejaswi D. Sudhakar, Ankita Saxena, Cole Chokran, Nelson Humala, Aayushi Mahajan, Weihao Xu, Jordan B. Metz, Cady Chen, Eric A. Bushong, Daniela Boassa, Mark H. Ellisman, Elizabeth M.C. Hillman, Guy M. McKhann, Brian J. A. Gill, Steven S. Rosenfeld, Catherine A. Schevon, Jeffrey N. Bruce, Peter A. Sims, Darcy S. Peterka, Peter Canoll

## Abstract

Gliomas are highly aggressive brain tumors characterized by poor prognosis and composed of diffusely infiltrating tumor cells that intermingle with non-neoplastic cells in the tumor microenvironment, including neurons. Neurons are increasingly appreciated as important reactive components of the glioma microenvironment, due to their role in causing hallmark glioma symptoms, such as cognitive deficits and seizures, as well as their potential ability to drive glioma progression. Separately, mTOR signaling has been shown to have pleiotropic effects in the brain tumor microenvironment, including regulation of neuronal hyperexcitability. However, the local cellular-level effects of mTOR inhibition on glioma-induced neuronal alterations are not well understood. Here we employed neuron-specific profiling of ribosome-bound mRNA via ‘RiboTag,’ morphometric analysis of dendritic spines, and in vivo calcium imaging, along with pharmacological mTOR inhibition to investigate the impact of glioma burden and mTOR inhibition on these neuronal alterations. The RiboTag analysis of tumor-associated excitatory neurons showed a downregulation of transcripts encoding excitatory and inhibitory postsynaptic proteins and dendritic spine development, and an upregulation of transcripts encoding cytoskeletal proteins involved in dendritic spine turnover. Light and electron microscopy of tumor-associated excitatory neurons demonstrated marked decreases in dendritic spine density. In vivo two-photon calcium imaging in tumor-associated excitatory neurons revealed progressive alterations in neuronal activity, both at the population and single-neuron level, throughout tumor growth. This in vivo calcium imaging also revealed altered stimulus-evoked somatic calcium events, with changes in event rate, size, and temporal alignment to stimulus, which was most pronounced in neurons with high-tumor burden. A single acute dose of AZD8055, a combined mTORC1/2 inhibitor, reversed the glioma-induced alterations on the excitatory neurons, including the alterations in ribosome-bound transcripts, dendritic spine density, and stimulus evoked responses seen by calcium imaging. These results point to mTOR-driven pathological plasticity in neurons at the infiltrative margin of glioma – manifested by alterations in ribosome-bound mRNA, dendritic spine density, and stimulus-evoked neuronal activity. Collectively, our work identifies the pathological changes that tumor-associated excitatory neurons experience as both hyperlocal and reversible under the influence of mTOR inhibition, providing a foundation for developing therapies targeting neuronal signaling in glioma.

## Introduction

Gliomas are the most common type of primary brain tumor in adults, who often present with neurological symptoms that include cognitive impairment, focal cortical dysfunction, and seizures. All these symptoms contribute to the progressive morbidity and mortality associated with this disease^1,2^. Gliomas grow by diffusely infiltrating the surrounding brain tissue, where they intermingle with non-neoplastic brain cells, including neurons^3^. Several studies have demonstrated that gliomas disrupt neuronal excitatory/inhibitory balance^4-8^ as well as functional remodeling of the peritumoral cortex^9-12^. Importantly, glioma-induced alterations of neuronal activity evolve as tumors progress, eventually leading to overt discharges and seizures in both mouse models^4,8,13,14^ and humans^15-18^. Notably, seizures and functional reorganization primarily occurs at the infiltrative margin of glioma^10,19-21^, and it has been proposed that glioma-associated neuroplasticity may contribute to the development of these seizures^1,12,22^. There is also evidence from experimental models that glioma growth and proliferation are affected by neuronal activity^21,23-33^, further highlighting the clinical importance of developing effective ways to normalize neuronal function as a means to treat neurological symptoms and slow tumor growth. However, glioma-induced neuronal alterations have yet to be adequately characterized on a cellular-level either by in vivo imaging, molecular profiling, or structural morphology.

Previous studies have shown that the glioma microenvironment contains elevated levels of mTOR activity and have implicated the consequently increased mTOR signaling in peritumoral neurons as contributing to a potentially targetable driver of glioma-associated hyperexcitability and seizures^8,34-36^. While mTOR hyperactivation, particularly in neurons, is believed to drive hyperexcitability through dysfunctional structural dynamics and receptor-level changes^37,38^, the exact role that mTOR signaling plays in the cellular-level glioma-induced neuronal alterations are not well understood. To address this, we have developed diffusely infiltrating glioma mouse models that are genetically engineered to allow for the detailed investigation of the phenotypic, morphologic, and functional changes in tumor-associated neurons. These mice often develop seizures during tumor progression, which we have previously characterized both by electrophysiological recordings and by in vivo GCaMP (calcium) imaging of neuronal activity^8,13^. Furthermore, we have shown that the mTORC1/2 inhibitor AZD8055 suppressed glioma-induced seizures in vivo, as well as the glioma-induced epileptiform activity in acute brain slices taken from the Thy1-GCaMP mouse glioma model^8^. However, the detailed in vivo cellular-level changes in tumor-associated excitatory neurons driven by the local effects of tumor-burden, and the effect of mTOR inhibition on these alterations are not well understood.

In this study, we used three complementary mouse models to assess the translational, morphological, and functional alterations of excitatory neurons in the infiltrated cortex, and the effects that AZD8055 has on these “tumor-associated neurons”. Transcriptional profiling of ribosome-bound mRNA within excitatory neurons in the CamK2A-RiboTag mouse glioma model^39^ revealed glioma-induced alterations of genes involved in synaptic function and dendritic spine plasticity, and that these gene expression alterations were reversed within six hours of AZD8055 treatment. Morphological analysis of tumor-associated excitatory neurons in Thy1-EGFP-M mice revealed a loss of dendritic spines, which was also reversed by AZD8055. Furthermore, in vivo two-photon calcium imaging in the Thy1-GCaMP6f mouse glioma model showed a significant increase in whisker stimulus evoked neuronal events, at multiple spatial scales, and increased neuronal synchrony as the tumors progressed from early to late stage. This imaging also revealed that these increases in stimulus evoked calcium events are the most pronounced in neurons that are in closest proximity to infiltrating glioma cells, and that these stimulus evoked events are suppressed by AZD8055. Additionally, tumor-associated neurons demonstrated alterations in temporal tuning to the whisker stimulus, which was also reversed following AZD8055 administration. Together, these findings show that neurons in the glioma infiltrated cortex undergo translational, morphological, and functional alterations, characterized by pathological plasticity and hyperexcitability and that these alterations can be reversed within hours of a single dose of AZD8055.

## Results

### Glioma-induced alterations in excitatory neuronal translation are reversed by AZD8055

To explore the molecular basis of neuronal alterations in glioma, we used a Camk2a-Cre driven RiboTag (Rpl22^HA^) transgenic mouse model, which allows for the isolation of transcripts bound to HA-tagged translating ribosomes specifically from excitatory neurons^39^. We implanted p53^-/-^ PDGFA^+^/mCherry-Luciferase^+^ murine glioma cells into the right frontal cortex of Camk2a-Cre-RiboTag mice. The mice developed diffusely infiltrating tumors with mCherry+ glioma cells intermingled with HA-tagged cortical excitatory neurons (**Fig. 1a**). We obtained RNA from glioma-infiltrated cortex, and we subsequently performed RNA sequencing on either total homogenate or from immunoprecipitated (IP) HA-tagged ribosomes (**Supplemental Figure 1a**). This enabled us to calculate the enrichment score of genes in the neuronal IP fraction and identified genes that were significantly enriched in neurons within the tumor brains (**Supplemental Table 1**). Gene set enrichment analysis (GSEA) using previously published neuron-specific gene signatures^40,41^ demonstrated that the mRNA from the IP polysome fraction was significantly enriched in neuronal genes (**Fig. 1b**, **and Supplemental Figure 1b**). Our approach is therefore suitable for probing neuron-derived genes from the tumor microenvironment, even though the fraction of neurons in the tumor is lower compared to the normal brain^13^ (**Supplemental Figure 1c**). Crucially, neuronal cell death, apoptosis, and autophagy pathways were not over-represented in the neuronal-enriched genes, indicating that our findings derive mainly from reactive, but not dying, neurons (**Supplemental Table 1**).

**Figure 1:**
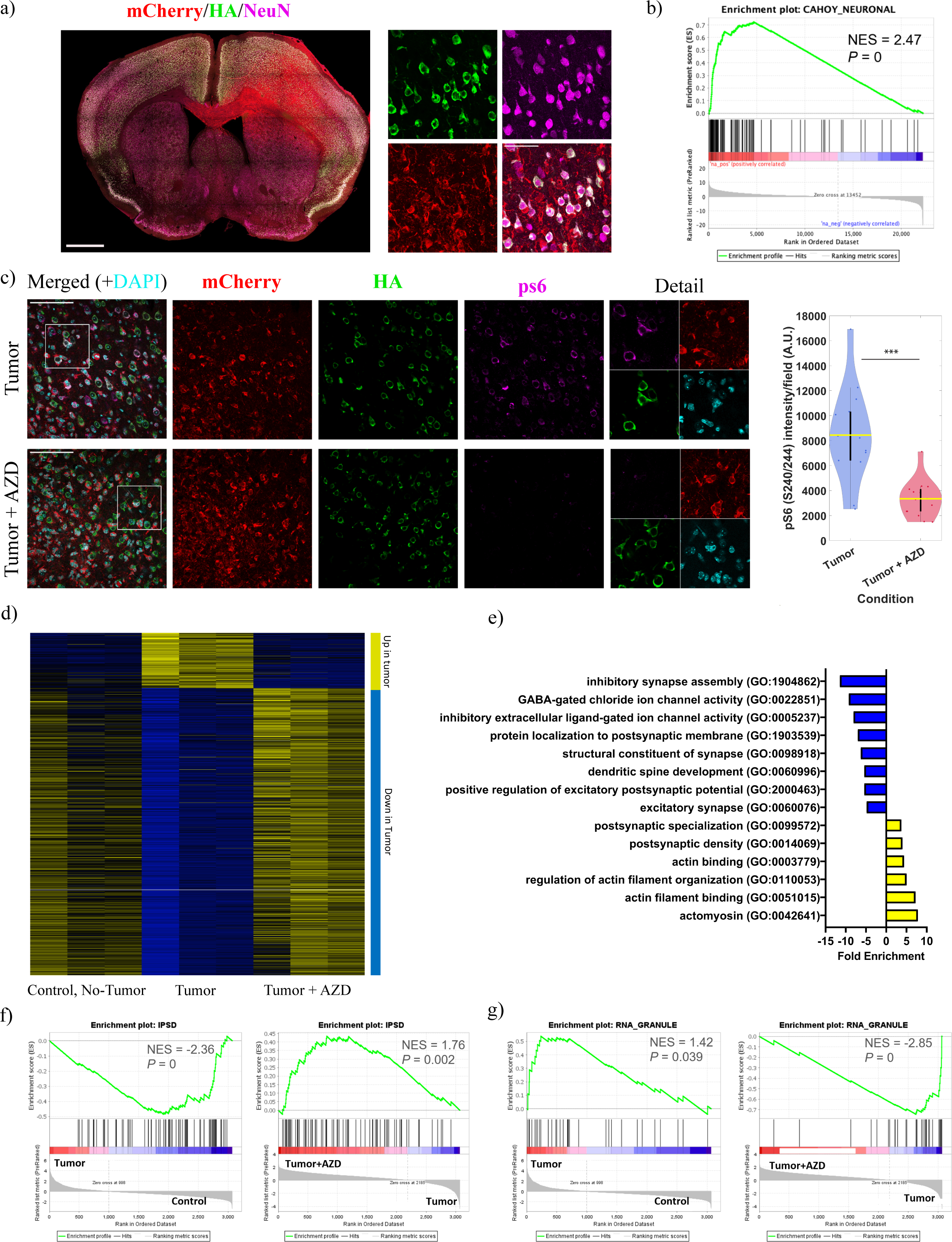
Tumor-Associated Neurons Show an mTOR-Dependent Translational Signature. a) Left: Coronal section of a mouse model of infiltrative glioma model showing an intermingling of mCherry+ glioma cells (red) with RiboTag (HA+; green) neurons. NeuN (magenta) is used to identify all neurons. Bar, 1mm. Right: Enlarged field from the cortex, showing prominent perineuronal satellitosis. Bar, 50 μm. b) GSEA plot showing enrichment of neuronal genes in the immunoprecipitated (IP) fraction of glioma-infiltrated brain. N = 3 mice per condition. c) Oral administration of AZD8055 effectively inhibits mTOR signaling in HA+ neurons in glioma-bearing mice. Representative images of cortex from mice treated with vehicle (top) or AZD8055 (bottom). Bar: 100 μm. Right: graph showing fluorescence intensity of pS6 (Ser240/244) levels in HA+ neurons, p=0.0004. N = 3 mice per condition, 15 image fields per condition. Wilcoxon rank-sum test was used. Yellow lines represent the mean. d) Map showing differential expression of neuronal-enriched genes between control brain, tumor brain, and tumor brain treated with AZD8055. N = 3 mice per condition. e) Gene ontologies that are altered between control and tumor brain and reversed by mTOR inhibition. N = 3 mice per condition. f) GSEA demonstrates depletion of inhibitory synapse genes in tumor brain compared to control (left) and an enrichment following AZD8055 treatment (right). N = 3 mice per condition. g) GSEA demonstrates enrichment of spine genes found in RNA granules in tumor brain compared to control (left) and their depletion following AZD8055 treatment (right). N = 3 mice per condition.

The mammalian target of rapamycin (mTOR) pathways play roles in both glioma progression^42-45^ and glioma associated epileptogenesis^8,34^. We have identified that acute ex vivo tumor-bearing slices from mice pre-treated with AZD8055^46^, exhibited reduced neuronal hyperexcitability^8^. A single oral dose of 100 mg/kg AZD8055 can rapidly reduce mTOR signaling within six hours in the naive brain (**Supplemental Figure 1d**). Administration of AZD8055 resulted in a marked reduction in phosphorylation of ribosomal protein S6 in tumor-associated HA+ neurons within six hours, showing that it can effectively inhibit mTOR signaling (**Fig 1c**). We performed RNAseq on IP fractions from the frontal cortex of control no-tumor brains, tumor-containing brains, and AZD8055-treated tumor-containing brains followed by DESeq2 analysis. We identified genes under the following three criteria: 1) neuronally enriched (significantly enriched in the IP fraction), 2) significantly differentially expressed between tumor-untreated and control-no-tumor brains, and 3) significantly differentially expressed between tumor-untreated and tumor-AZD8055-treated brains. DESeq2 analysis identified both up- and downregulated gene expression that depended on the gene and the comparison (**Supplemental Figure 1e, Supplementary Table 1**). Remarkably, comparing these gene lists revealed that a statistically significant majority of the neuron-enriched genes downregulated in tumor were upregulated following AZD8055 treatment, while those upregulated in tumor were downregulated following AZD8055 (**Fig 1d, Supplemental Figure 1e, Supplementary Table 1**). We subsequently focused our analysis on these genes that met the above-mentioned three criteria.

Gene ontology (GO) analysis of the neuron-enriched genes that were downregulated in tumor and upregulated following AZD8055 revealed involvement in inhibitory synapse assembly, GABA-gated ion channel activity (e.g., Gabra1, Gabrb2, Gabrg2), and other ontologies associated with dendritic spines and synapses (**Fig 1e; blue bars**). This is consistent with the loss of neuropil elements in the tumor infiltrated cortex, as identified by a neuropil specific geneset^47^ (**Supplemental Figure 1f**). Next, to further explore the effects of glioma-induced alterations and mTOR inhibition on excitatory and inhibitory synapses, we performed GSEA analysis on gene sets associated with excitatory and inhibitory post-synaptic densities^48^ (**Supplementary Table 1**). This analysis revealed that both inhibitory (**Fig. 1f**) and excitatory post-synaptic genes (**Supplemental Figure 1f**) are significantly downregulated in tumor and significantly upregulated following AZD8055.

By contrast, GO analysis revealed that a subset of neuron-enriched genes encoding for cytoskeletal proteins was upregulated in tumor and downregulated following AZD8055. (**Fig. 1e; yellow bars, Supplemental Table 1**). Previous studies have shown that many of these genes are involved in dendritic spine turnover^49,50^, and that their transcripts are enriched in RNA granules associated with local translation near dendritic spines^51^. Notably, GSEA revealed that the RNA granule signature from El Fatimy et al.^51^ (**Supplemental Table 1**), is significantly enriched in the glioma-associated neurons, upregulated in tumor, and downregulated following AZD8055 treatment (**Fig 1g**). Furthermore, as noted above, the specific GO of dendritic spine development is significantly downregulated in the tumor. Taken together, an elevation in genes responsible for dendritic spine turnover and a reduction in genes responsible for dendritic spine development suggests a net loss of dendritic spines in tumor-associated excitatory neurons. Therefore, we sought to directly measure the levels of dendritic spines and inhibitory postsynaptic structures in glioma-associated excitatory neurons both pre- and post-AZD8055.

### Reversible changes in dendritic spine and inhibitory synapse densities

Dendritic spines are highly dynamic structures that undergo actomyosin-dependent changes in density^52,53^. To directly assess the effect of gliomas on dendritic spine morphology and density in excitatory neurons and the effects of mTOR inhibition, we performed DiOlistic labeling of slices from the Cank2a-RiboTag mouse model including control no-tumor brains, tumor-containing brains, and AZD8055-treated tumor-containing brains and we counted dendritic spines from HA^+^ neurons that were labeled with DiI. Olig2 was used as a marker to determine the presence of tumor in each field (**Fig 2a**). We observed a 3-fold decrease in the density of dendritic spines in glioma-associated excitatory neurons compared to excitatory neurons in control no-tumor brains. Serial scanning electron microscopy and reconstruction of dendrite segments at the margins of the glioma demonstrated a similar decrease in spine density (**Supplemental Figure 2a, b**). Remarkably, within six hours of a single oral dose of AZD8055 (100 mg/kg), the dendritic spine density was partially restored (**Fig 2a, Supplemental Figure 2a, b**). This suggests that the alterations in dendritic spines induced by glioma infiltration are reversible and regulated by mTOR signaling.

**Figure 2:**
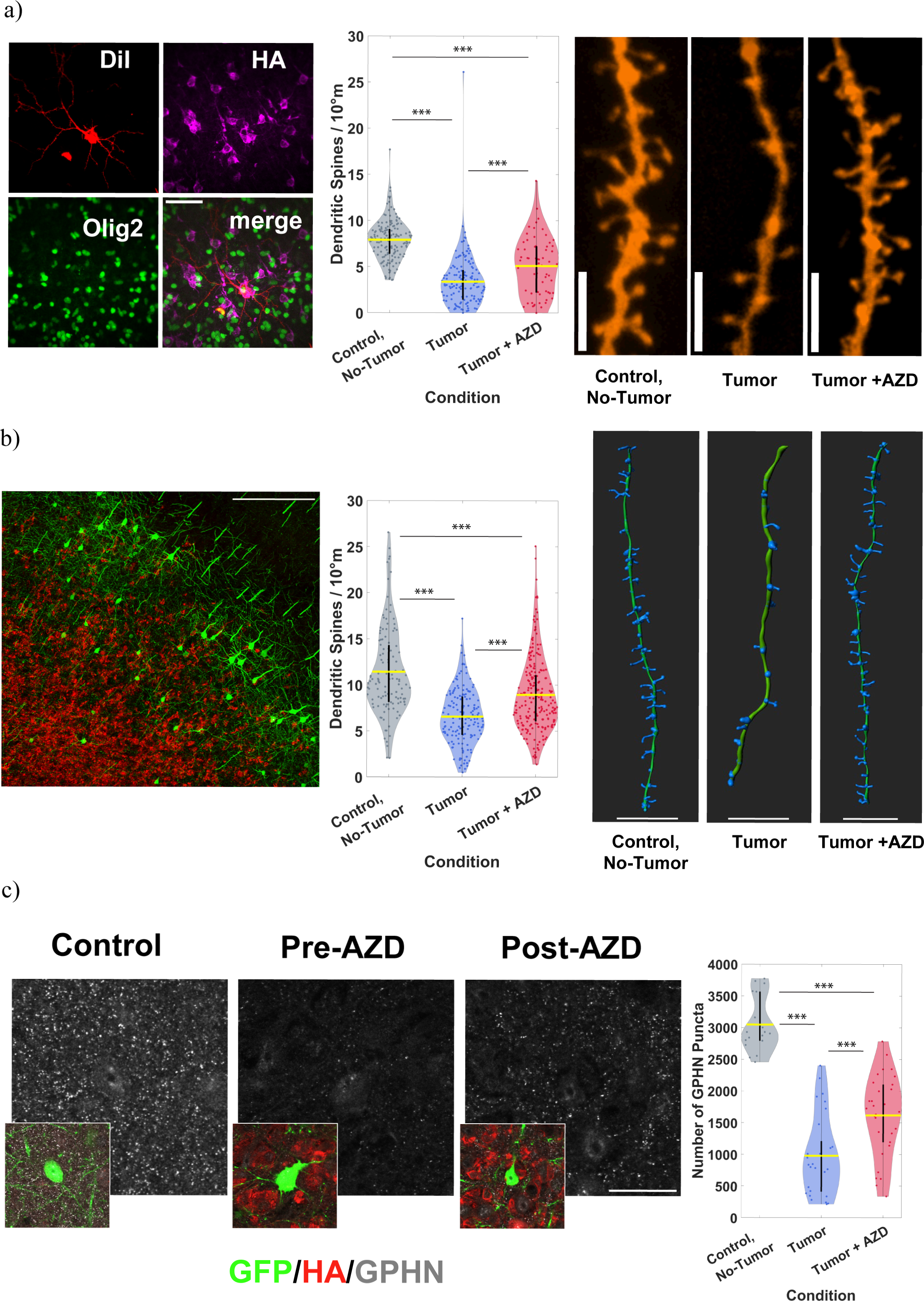
mTOR-dependent Alterations in Dendritic Spine and Inhibitory Synapses. a) Left: representative field from a DiI-labeled (red), HA+ neuron (magenta) in Olig2+ (green) glioma-infiltrated cortex. Scale Bar: 50 μm. Graph: violin plot of dendritic spine densities among conditions. Right: representative images of dendrites. Scale Bar: 5 μm. Control no-tumor vs. tumor (p =1.8197e-38). Control vs. tumor+AZD8055 (p = 3.0766e-10). Tumor vs. tumor+AZD8055 (p = 6.8757e-05). N = 155 dendrites across 5 control mice, 152 dendrites across 9 tumor mice, 64 dendrites across 5 tumor+AZD8055 mice. Wilcoxon rank-sum test was used. Yellow lines represent the mean. b) Left: representative field of the glioma-infiltrated cortex (glioma cells: mCherry; red) of a Thy1-EGFP- M mouse (neurons; green). Bar: 250 μm. Graph: violin plot of dendritic spine densities among conditions. Right: representative Imaris-rendered images of dendrites. Scale Bars: 10 μm. Green: dendrite, blue: dendritic spines. Control vs. tumor (p = 4.6216e-20). Control vs. tumor+AZD8055 (p = 3.2152e-07). tumor vs. tumor+AZD8055 (p = 8.5802e-09). N = 123 dendrites across 4 control mice, 161 dendrites across 3 tumor mice, 222 dendrites across 5 tumor+AZD8055 mice. Wilcoxon rank-sum test was used. Yellow lines represent the mean. c) Left: Representative confocal sections of the cortex of Thy-EGFP-M mice showing gephyrin puncta (gray). Scale Bar, 25 μm. Insets show GFP+ neurons and mCherry+ glioma cells in the same field. Right: violin plot showing changes in gephyrin puncta per field. Control vs. tumor (p = 1.1919e-08). Control vs. tumor+AZD8055 (p = 2.2665e-08). tumor vs. tumor+AZD8055 (p = 5.9423e-04). N = 18 imaging fields across 3 control mice, 29 imaging fields across 4 tumor mice, 30 imaging fields across 4 tumor+AZD8055 mice. Wilcoxon rank-sum test was used. Yellow lines represent the mean.

To validate this finding, we implanted the mouse glioma cells into a) the frontal cortex and b) the motor cortex of Thy1-EGFP-M mice, in which excitatory cortical neurons are sparsely labeled with EGFP. In our analysis of glioma-associated excitatory neurons at the infiltrated margins in both the frontal cortex (**Fig 2b**) and the motor cortex (**Supplemental Figure 3a**), we again found a similar glioma-induced decrease in dendritic spine density, and significant increases in dendritic spines following AZD8055.

This glioma-induced loss of dendritic spines is consistent with our results seen in the RiboTag analysis – specifically, the significant down-regulation of both excitatory postsynaptic assembly genes and dendritic spine development GOs, as well as the significant upregulation of cytoskeletal GOs containing dendritic spine turnover genes (**Fig 1**, **Supplemental Table 1**). Given this apparent loss of excitatory input, the hyperexcitability we have observed in this model^8,13^ may seem surprising. However, our RiboTag data also revealed a significant decrease in several transcripts associated with the postsynaptic compartment of inhibitory (GABAergic) synapses in excitatory neurons (**Fig 1f, Supplemental Table 1**). Since the loss of local inhibitory input onto excitatory neurons has been proposed to contribute to the neuronal hyperexcitability and seizures that are commonly seen clinically in patients with glioma^7,8,14,54^, we stained brain sections of the Thy1-EGFP-M mice with an antibody against gephyrin (GPHN), a scaffold protein important in the assembly of signaling and structural components of postsynaptic apparatus of inhibitory synapses^55^. Compared to control brains, we found that tumor-bearing brains show a dramatic decrease in the number of GPHN+ puncta, suggesting a decrease in inhibitory synapses. Importantly, the number of total GPHN+ puncta in the infiltrated field (**Fig 2c**) and the number of GPHN+ puncta localized to GFP^+^ neurons (**Supplemental Figure 2b**) was significantly increased following AZD8055 administration, though GPHN puncta remained lower than that of control no-tumor brains. These findings show that mTOR inhibition can reverse the glioma-induced loss of dendritic spines and inhibitory synapses in excitatory neurons. Next, we sought to directly assess the functional consequences of these effects.

### In vivo two-photon imaging reveals progressive alterations in tumor-associated excitatory neurons

Several groups have utilized calcium imaging to evaluate glioma-induced changes in neuronal activity^13,14,56^. To follow alterations in neuronal activity through the process of tumor progression, we implanted tumor cells adjacent to the somatosensory cortex and performed longitudinal two-photon imaging of excitatory neurons in the whisker barrel cortex of awake Thy1-GCaMP6f^57^ mice and recorded the responses to contralateral whisker stimulation (**Fig 3a**). Tumor cells progressively invaded the whisker barrel cortex (**Fig 3b**), and we performed longitudinal imaging of calcium activity in tumor-associated excitatory neurons at the infiltrative edge of the tumor in this cortical region. Calcium events were recorded and analyzed at the level of the entire imaging field-of-view (**Supplemental Movie 1**), as well as at the level of the individual neuron (**Supplemental Movie 2**). Tumor imaging fields always had more field-level events during 3- minute whisker stimulation runs than did fields from control no-tumor brain, regardless of event threshold (**Supplemental Figure 3b**). Moreover, we observed events with very high amplitudes in the tumor data, and such large events never occurred in the control no-tumor animals. A subset of these high amplitude events were also synchronous with the onset of whisker stimulation. These large field-level events typically lasted short durations (<1s) and were found to originate at the invasive tumor margin (**Fig 3c, 3d; Supplemental Movie 1)**. To quantify the progressive occurrence of high-amplitude field-level calcium events during whisker stimulation runs, we calculated the mean ΔF/F and standard deviation across each imaging field-of-view and set the threshold for events at five standard deviations above the mean signal. We found that these high amplitude field-level calcium events significantly increased as tumors progressed from early (<21 DPI) to late (>21 DPI) (**Fig 3e**). We recognized that these field-level events bore a striking resemblance to data presented by Liu et al. (2021)^58^ in a mouse model of STXBP1 associated epilepsy, in which “cascade” events were characterized as transient periods of elevated activity among spatiotemporally contiguous pixels from GCaMP recordings. We applied established methods from prior studies^58-61^ to detect cascades in our raw data and found that these field-level events appeared as cascades increasing in both spatial size and duration as the tumors progressed (**Supplemental Figure 3c**).

**Figure 3:**
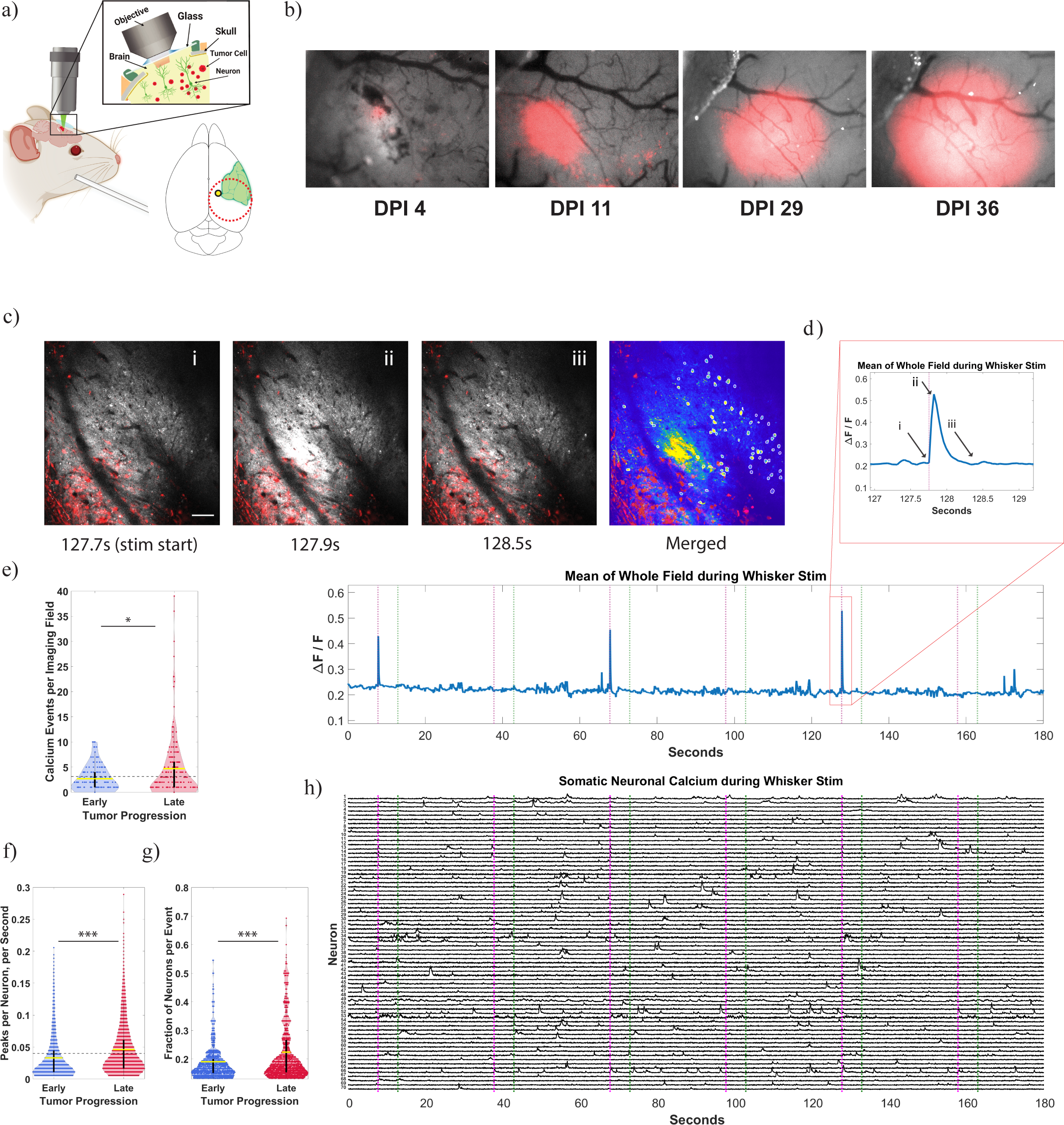
In Vivo Two-Photon Imaging Reveals Progressive Alterations in Tumor-Associated Excitatory Neurons. a) Left: Schematic of in vivo head-fixed two-photon imaging with simultaneous whisker stimulation (modified from Hillman, 2007). Right: Schematic of tumor injection placement (yellow marker) with respect to somatosensory cortex (green area, modified from Kirkcaldie et al., 2012). b) Representative low magnification images of tumor cell fluorescence in the cranial window growing over the course of tumor progression. mCherry-tagged tumor cells were excited by 590nm LED and the resulting signal was pseudo colored to red in the figure. c) Representative images of a large amplitude field-level event occurring during whisker stimulus onset, in an imaging field of a tumor bearing animal at 29 DPI. Three different still frames (i, ii, iii), with calcium signal from GCaMP in grayscale and maximum intensity projection from a 200um z-stack of the tumor in red, show the close temporal alignment with stimulus onset, the sharp rise of the event, and short duration. The timeseries of this event is plotted in figure 3d. The merged image of the same structural maximum intensity projection of the tumor in red overlaid with a maximum intensity projection of the calcium signal through time of the three-minute whisker stimulation run, shows the large amplitude field-level events located at the tumor margin as well as the somatic neuronal ROIs. Scale bar is 100um. d) Larger timeseries below shows calcium ΔF/F during the entire three-minute imaging run generated from the mean of the whole imaging field. The pink vertical lines represent the start of each whisker stimulation epoch, and the green vertical lines represent the end of each whisker stimulation epoch, separated by 25-second no-stim intervals. During this run, there were three large amplitude field-level events associated with stimulus onsets, with increasing amplitude. The last and largest event is plotted again on the zoomed inset, with the timing of each frame from 3c marked (i, ii, iii). Two-photon imaging was performed at 30Hz. e) Mean ΔF/F and standard deviation across each imaging field-of-view (512×512 pixels) during three-minute whisker stimulation runs were recorded and thresholds were set for calcium events at five standard deviations above the mean signal. The number of calcium events above these thresholds were significantly higher (p=0.0170) in recordings from late tumor progression animals (>21 DPI, N=182 recordings across 7 mice) than recordings in early progression animals (<21 DPI, N=137 recordings across 7 mice). Wilcoxon rank-sum test was used. Yellow lines represent the mean. The dotted black line represents the mean number of discharges per imaging field in the control no-tumor recordings (N=124 recordings across 5 mice). f) Neuronal somatic calcium events per second were computed during three-minute whisker stimulation runs. Mean ΔF/F and standard deviation for each neuron was calculated and thresholds were set for calcium events at six standard deviations above the mean signal. Significant differences (p = 3.7221e-111) were observed when comparing the event rate of all neurons from early DPI recordings (N=4142 neuronal ROIs across 6 mice) to the event rate of all neurons from late DPI recordings (N=6622 neuronal ROIs across 7 mice). Wilcoxon rank-sum test was used. Yellow lines represent the mean. The dotted black line represents the mean event rate across the control no-tumor neurons (N= 9978 neuronal ROIs across 5 mice). g) A synchronous event was identified as at least two different neurons in a field-of-view having a calcium peak occurring in a 500-millisecond interval during a five-second whisker stimulation epoch. Displayed are the fraction of neurons involved per synchronous event, restricted to events that contained at least∼13% of neuron involvement (two standard deviations above the mean of the control no-tumor data, N= 2340 synchronous events across 5 control mice). The fraction of neurons involved per synchronous event was significantly higher (p = 3.2086e-25) in FOVs from later DPI recordings (N= 3272 synchronous events across 7 mice) than from early DPI recordings (N= 2583 synchronous events across 6 mice). Wilcoxon rank-sum test was used. Yellow lines represent the mean. h) Corresponding calcium ΔF/F time series from all the neuronal somatic ROIs in the field of view from 3c, aligned to the same whisker stimulation, illustrate that large field-level calcium measurements are a distinct and complementary measurement to single neuron-level measurements.

Next, using established methods^62^, we create regions of interest (ROIs) around neuronal somata and extracted the ΔF/F from the these neuronal ROIs to analyze calcium events on the single-neuron level (**Fig 3c, 3h**). Interestingly, these large field-level events were not seen in the signals from neuronal somata, demonstrating that this field-level calcium analysis provides a distinct measurement of neuronal activity that is complementary to the single-neuron level measurement. During the full three-minute whisker stimulation runs, single-neuron calcium events were significantly increased in late-tumor mice when compared against early-tumor mice (**Fig 3f**). Increased neuronal synchrony has been identified as a central pathological component of seizures and seizure progression^63-65^. We used an established method^63^, with slight modifications, to identify synchronous events within an imaging field as any 500-millisecond interval where two or more neurons had a calcium event above threshold. We found that the fraction of neurons in an imaging field that participated in each synchronous event during five-second whisker stimulation epochs increased during tumor progression (**Fig 3g**). Neuronal recordings from both early and late tumors exhibited greater neuronal recruitment in synchronous events than control no-tumor animals (**Supplemental Figure 3d**).

### In vivo two-photon imaging reveals the modulatory effects of mTOR inhibition on excitatory neuronal activity at the infiltrative glioma margin

Next, to evaluate the effects of mTOR inhibition on these alterations in neuronal activity, AZD8055 was administered to five mice on the last day of their longitudinal imaging (**Fig 4a, Supplemental Table 2**), and imaging was performed before and after AZD8055 administration. Prior to AZD8055, the fraction of neurons involved in synchronous events during whisker stimulation epochs was significantly higher than control no-tumor animals. After administration of AZD8055 the fraction of neurons involved in each synchronous event was significantly reduced compared to the pre-treatment condition (**Fig 4b**). Consistent with previous observations^66,67^, we saw neurons that responded during periods of whisker stimulation (**Fig 4c**). Additionally, we noted that at any time point along tumor progression throughout the infiltrative tumor margin, some neurons were in close proximity to tumor cells while others were farther from tumor cells (**Fig 4d**). Therefore, to analyze the local effects of tumor cells on calcium activity on a neuron specific basis, we used structural image stacks to compute a tumor-burden score for each neuronal ROI based on its proximity to tumor cells in the microscopic field and then grouped neuronal ROIs as high- or low-tumor-burden (**Fig 4d**, **Supplemental Figure 5a**). We observed that the high-tumor-burdened neurons have particularly elevated calcium peak rates during the full three-minute whisker stimulation runs relative to low-tumor-burden or control no-tumor neurons, and that this elevated event rate was reduced to control levels following AZD8055 administration (**Fig 4e)**. To further quantify this effect, we looked at the rate of neuronal calcium events above threshold specifically during periods of whisker stimulation (five-second whisker stimulation epoch plus two-seconds after). We found that before AZD8055 administration, the high-tumor-burden neurons had significantly elevated calcium event rates during these whisker stimulation periods relative to control no-tumor neurons as well as low-tumor-burden neurons (**Fig 4c and 4f**). Following AZD8055 administration, the calcium event rates of the high-tumor burden neurons, low-tumor-burden neurons, and control no-tumor neurons no longer statistically differed during whisker stimulation periods.

**Figure 4:**
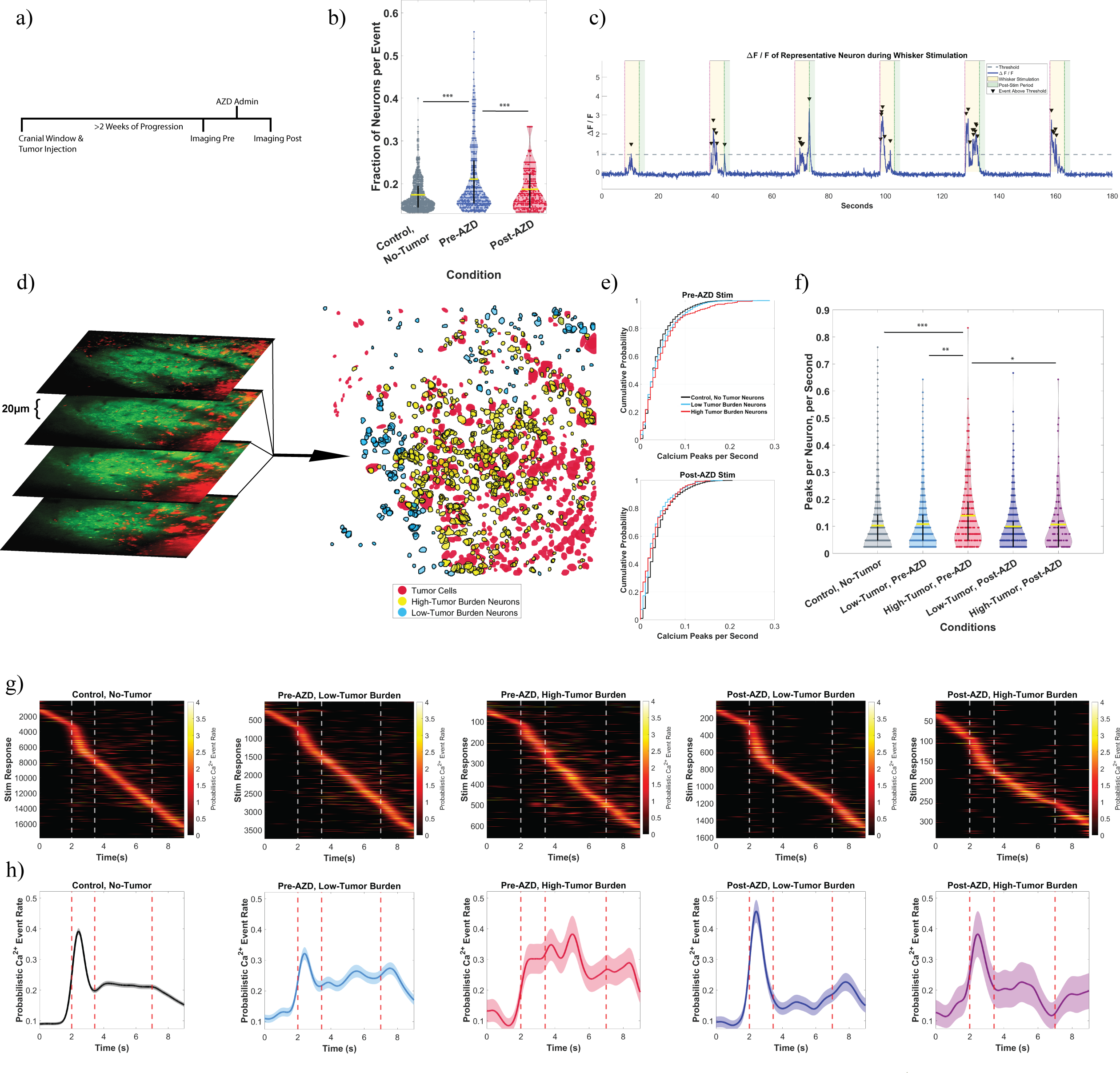
In vivo Two-Photon Imaging Reveals the Modulatory Effects of mTOR Inhibition on Excitatory Neuronal Activity at the Infiltrative Glioma Margin. a) Representative timeline of experimental design to record functional in vivo alterations in tumor-associated neurons in awake animals using two-photon imaging before and after mTOR inhibition via AZD8055. b) The fraction neurons involved per synchronous event was significantly higher between the tumor-associated neurons pre-AZD8055 and control no-tumor neurons (p = 1.6733e-29) and was significantly lowered after AZD8055 administration (p = 8.8852e-08). The post-AZD8055 group remained significantly higher than the control no-tumor (p = 9.0423e-06). N = 5 mice in each condition. Wilcoxon rank-sum test was used. Yellow lines represent the mean. c) Representative time series of ΔF/F fluorescence from a single stimulus-tuned neuronal ROI during whisker stimulation protocol, with peaks above the threshold indicated. Each whisker stimulation run was three-minutes, including six five-second whisker stimulation epochs separated by 25-second interstimulus intervals. d) Left: Structural imaging of tumor cells was overlaid with maximum intensity projections of functional neuronal calcium to determine the tumor burden of each neuron (red is tumor signal, and green is neuron calcium signal). Right: Representative image of a field of view with tumor cell regions of interest (red), low-tumor-burden neuron regions of interest (blue), and high-tumor-burden neuron regions of interest (yellow). See Supplemental Figure 5a for further details on methodology. e) Cumulative probability plots of neuronal calcium peaks per second, during the full three-minute calcium imaging runs, illustrate that the largest observable difference in the calcium peak rate is found in the increased peaks per minute of the high-tumor-burden neurons, during whisker stimulation, before AZD8055 administration (top panel). These differences were not observed following AZD8055 administration. f) Neuronal somatic calcium events per second were computed during whisker stimulation periods –the five-second stim epoch plus two seconds immediately following whisker stimulation. Differences were observed between high-tumor pre-AZD8055 and the control no-tumor (p = 2.0795e-05), high-tumor pre-AZD8055 and low-tumor, pre-AZD8055 (p = 0.0011). After AZD8055 treatment, there were no significant differences between high-tumor post-AZD8055 and control no-tumor (p = 0.8180), low-tumor post-AZD8055 and control no-tumor (p = 0.6438), high-tumor post-AZD8055 and low-tumor post-AZD8055 (p = 0.9975). N = 9978 neuronal ROIs across 5 control mice, 2226 pre-AZD8055 low tumor burden neuronal ROIs across 5 mice, 373 pre-AZD8055 high tumor burden neuronal ROIs across 5 mice, 1120 post-AZD8055 low tumor burden neuronal ROIs across 5 mice, and 277 post-AZD8055 high tumor burden neuronal ROIs across 4 mice. Wilcoxon rank-sum test was used. Yellow lines represent the mean. g) Plotted are all the individual temporally smoothed stimulus evoked calcium responses across all neurons in each condition. The responses are sorted according to the timing of their maximum response, from earliest to latest. The first and last vertical dashed lines indicate stimulus onset and offset. The middle vertical dashed line indicates the position of the end of the average control response to the stimulation onset. N = 17879 control stimulation responses across 5 mice, 3741 pre-AZD8055 low tumor burden stim responses across 5 mice, 659 high tumor burden stim responses across 5 mice, 1611 post-AZD8055 low tumor burden stim responses across 5 mice, 342 post-AZD8055 high tumor burden stim responses across 4 mice. h) Plotted are the average stimulus evoked responses for each condition, across all animals, from the corresponding data in Fig 4g.

To better understand the temporal relationship between the activity of the various groups of neurons and the whisker stimulation, the neuronal time series were aligned within each group (**Supplemental Figure 5b**). Following temporal smoothing of the calcium peaks (**Supplemental Figure 4**), average traces of calcium peaks per second were computed for each group of neurons during whisker stimulation (**Fig 4g, 4h**). The averaged trace for the control no-tumor neurons showed a consistent patterned stimulus-evoked response characterized by a prominent peak of highest average response during the first second of stimulation followed by a plateau of less elevated response during the remaining four seconds of stimulation, then returning toward baseline. This pattern was disrupted in both low- and high-tumor-burden neurons and was significantly corrected following AZD8055 to varying degrees in each neuronal group. Administration of AZD8055 significantly increased the early component and decreased the later component of the responses in the low-tumor-burden neurons, and significantly decreased the later component of the responses in the high-tumor-burden neurons (**Supplemental Figure 5c**). Following AZD8055 administration, the stimulus-evoked responses of both the low- and high-tumor-burden neurons became more similar to that of the average control no-tumor response. When comparing both correlation coefficients and mutual information between the average low-tumor burdened neuron trace and the control no-tumor neuron trace, the correlation coefficients increased from R = 0.83 to R = 0.96 and the mutual information increased from 0.70 to 0.77 after AZD8055. The results when comparing the average high-tumor burdened neuron trace to the control no-tumor trace were more dramatic: the correlation coefficients increased from R = ™0.12 to R = 0.87 and the mutual information increased from 0.42 to 0.72 after AZD8055.

## Discussion

### Local neuronal alterations lead to hyperexcitability at the infiltrated margin of glioma

Gliomas are not only an aggressive form of brain cancer, but also cause devastating neurological disease, marked by cognitive dysfunction, language and memory impairment, and seizures. Activity alterations in the glioma associated neurons can eventually lead to interictal discharges and seizures, which often originate at or beyond the infiltrative margin of the tumor^68-71^. Cellular changes occurring at the infiltrative margin are likely caused by the complex multicellular spatial, chemical, and temporal interactions among the closely associated cells of the tumor microenvironment^3,72^. For example, in the glioma infiltrated cortex, glioma cells surround the cell bodies of excitatory pyramidal neurons forming a histopathological pattern referred to as ‘perineuronal satellitosis’. Glioma cells also diffusely infiltrate the cortical neuropil where they intermingle with neuronal processes, including axons and dendrites, establishing cellular interactions, with the potential to have hyperlocal effects on neuronal function^3,9^.

Our results show that tumor-associated neurons undergo pathological alterations in protein expression, morphology, and function in response to tumor infiltration. Furthermore, we show that the functional alterations of excitatory neurons become progressively more severe as more glioma cells infiltrate the whisker barrel cortex, including increased whisker stimulus evoked field-level calcium events, increased recruitment of neurons to synchronous events, and higher event rates.

Computing a tumor-burden score based on the local proximity of excitatory neurons to tumor cells revealed significant differences in the behavior of neurons that are within the same microscopic field, with more severe alterations seen in the high-tumor-burden neurons. Recent work in the field has identified synaptic interactions, such as AMPAR mediated synapses, and secreted factors, such as Glutamate, NLGN-3 and BDNF, as mediating crosstalk between neurons and glioma cells^21,23,29,31^. While most of these prior studies have focused on the effects of these interactions on the glioma cells, our analysis focused on glioma-induced alterations in excitatory neurons and revealed a loss of both inhibitory and excitatory post-synaptic proteins, as well as a loss of dendritic spines. The net consequence of the changes we observed is progressive cortical dysfunction and hyperexcitability that culminates in epileptiform activity. These findings of increasing hyperexcitability during glioma progression are consistent with those of previous studies^13,14^, and further highlight the role of local alterations at the level of individual neurons in generating seizures at the infiltrative margin of glioma.

### Physiologic Stimulus-Evoked Responses Reveal Glioma Associated Cortical Dysfunction

Glioma not only causes hyperexcitability and seizures, but can also alter task-based responses in functionally-connected regions of glioma-infiltrated cortex^10^. Furthermore, recent work has suggested that physiologic-level neuronal stimulation can influence glioma cell proliferation^24,28^, further highlighting the bi-directional interplay between brain function and glioma cells. Here we focused on functionally responsive somatosensory cortex at the infiltrative margin of glioma to understand how physiologic sensory stimulation can drive abnormal responses in tumor associated neurons. While some of the measurements of stimulus-evoked events were dramatic, inducing field-level calcium bursts and increased neuronal recruitment into progressively larger synchronous events, other alterations were more subtle – increases in overall event rate and disturbed average responses to stimulus timing. Again, the alterations in stimulus-evoked responses varied significantly across neurons, with high-tumor-burden neurons showing more severe alterations in event rate and larger distortion of the temporal alignment of the associated activity with the stimulus. This illustrates that even functionally responsive tumor-associated cortex displays subtle changes in stimulus-evoked responses which could progress to more severe cortical dysfunction. These functional alterations, along with the translational alterations in genes associated with synaptic plasticity, and the structural alterations seen in dendritic spines, point to neuronal plasticity as a key mechanism underlying glioma-induced cortical dysfunction, and all appear partially reversible by mTOR inhibition. Clinically, in humans, we know that pathological plasticity exists on a spectrum of severity in glioma patients, ranging from functional remodeling of neural circuits seen during intraoperative mapping^10,73,12^, to deficits in attention, concentration, processing speed, learning, memory, language, executive functioning, and seizures^74,75^. Our findings suggest that inhibiting mTOR could alleviate these devastating neurological symptoms.

### mTOR Signaling is a Master Regulator of Glioma-Induced Neuronal Alterations

Alterations in mTOR signaling have been implicated in a variety of neurological diseases with pathological mechanisms relating to mTOR’s critical role in local protein synthesis, dendritic spine morphogenesis and synaptic plasticity^76-80^. Inhibiting mTOR can reduce seizures in animal models of TSC-related epilepsy^81,82^ and can reverse TSC-associated alterations in synaptic plasticity with improvements in cognitive deficits^83^. Our findings show that inhibiting mTOR reverses glioma-induced alterations on neuronal translation, dendritic spine loss, neuronal hyperexcitability and response to whisker stimulation. Thus, mTOR provides a unifying mechanistic/therapeutic link between these different aspects of glioma-induced neuronal dysfunction. Of note, these changes occurred within just six hours of AZD8055 treatment, highlighting the dynamic reversibility of these glioma-induced neuronal alterations. As the gene expression differences were identified at the level of ribosomal associated mRNA, using the RiboTag IP, our results indicate that these neuronal expression changes are mediated at the level of translation, and likely relate to mTOR’s established role in regulating local translation in dendrites^84,85,86^. Previous studies have shown that a specific population of RNA granules are transported into the dendrite and localize at dendritic spines, and these granules contain transcripts of proteins involved in cytoskeleton remodeling, dendritic spine morphogenesis and synaptic plasticity^51^. Remarkably, we found that this dendrite spine-associated RNA granule gene signature was significantly enriched in the glioma-associated neurons compared to neurons from control no-tumor brains, and was significantly depleted following AZD8055 treatment, implicating mTOR signaling in both glioma-induced translational alterations and glioma-induced spine alterations. This was further supported by our morphological analysis, which showed a significant loss of dendritic spines in glioma-associated neurons that was reversed by AZD8055. Previous studies have shown that inhibiting mTOR signaling can induce increases or decreases in dendritic spine density, depending on the specific disease context^87-90^. One unifying explanation for these complex effects is that increased mTOR signaling drives pathological plasticity, and therefore inhibiting mTOR normalizes against the neurological insult, towards homeostasis, regardless of the direction of pathological alterations. Consistent with this idea, our findings show that neurons in the glioma infiltrated cortex undergo transcriptional, morphological, and functional alterations, characterized by pathological plasticity and hyperexcitability and that all of these alterations are significantly reversed within six hours of a single dose of AZD8055.

### Therapeutic implications of Reversibility of Neurological Alterations

Previously, we showed that inhibiting mTOR with AZD8055 increased the firing rate of putative fast-spiking interneurons and restored the GABA reversal potential in excitatory neurons, which increased inhibitory tone, and suppressed discharges and seizures^8^. The current study complements these findings by showing that AZD8055 also reverses the glioma-induced translational alterations of inhibitory postsynaptic genes expressed in excitatory neurons, thus restoring the ‘inhibitability’ of excitatory neurons. Together these findings suggest that inhibiting mTOR may reduce the neuronal alterations and cortical dysfunction that leads to cognitive impairment and seizure induced by gliomas. Our findings also identify new potential therapeutic strategies to slow glioma progression, by targeting mTOR-mediated neuronal hyperexcitability at the tumor margin, which would complement other treatments aimed at targeting neuron-glioma crosstalk. For example, our results suggest that AZD8055 will both reduce hyperexcitability and render excitatory neurons more sensitive to drugs that modulate inhibitory tone, such as traditional GABAergic anticonvulsants. These combined effects could improve glioma-associated seizure treatment, and other neurological dysfunctions, while also potentially reducing the growth-promoting effects that neuronal activity appears to have on glioma cells^21,23-33^. Furthermore, the loss of dendritic spines and postsynaptic structures that is seen at the margin of glioma may provide fertile ground for infiltrating glioma cells to successfully compete for synaptic input. AZD8055 or other drugs that reverse these pathological changes could shift the competitive balance within the highly plastic margin of glioma towards more normative function.

## Acknowledgments

We thank the Herbert Irving Cancer Center Confocal Core and the Zuckerman Institute’s Cellular Imaging core for instrument use and technical advice.

## Contributions

<u>Conceptualization:</u> A.R.G., A.D., D.T., P.A.S., P. Can.; <u>Methodology:</u> A.R.G., A.D., D.T., S.D.S, A. Me., E.M.M., M.O., L.A.S., H.T.Z., P. Can., M.A.B., W.X., J.B.M., E.A.B., D.B., M.H.E., E.M.C.H., B.J.A.G., C.A.S., P.A.S., D.S.P., P. Can.; <u>Investigation:</u> A.R.G., A.D., D.T., S.D.S, A. Me., M.O., C.K., A.V., A.R., T.D.S., A.S., C. Cho., N.H., A. Ma., J.B.M., C. Che., E.A.B., D.B.; <u>Writing - original draft:</u> A.R.G, A.D., D.T., P. Can.; <u>Writing - review & editing:</u> A.R.G., A.D., S.S.R., D.P., P. Can.; <u>Resources:</u> M.H.E., E.M.C.H., G.M.M., B.J.A.G., S.S.R., C.A.S., J.N.B., P.A.S., D.S.P., P. Can.; <u>Funding acquisition:</u> A.R.G., M.H.E., S.S.R., J.N.B., P.A.S., P. Can.; <u>Supervision:</u> M.H.E., C.A.S., P.A.S., D.S.P., P. Can.

## Methods

### Ethics statement

All experiments involving mice were approved by the Institutional Animal Care and Use Committee (IACUC) of Columbia University and performed in accordance with institutional policies.

### Tumor cell lines

The glioma cells used in this study have been previously described^13,91^. Glioma cells injected into Camk2a-Cre-RiboTag mice lacked the RPL22-HA transgene (APCL cell line). For in vivo imaging, the 333 cell line, which expressed RPL22-HA^13^, was infected with mCherry-expressing retrovirus to further enhance its excitation by two-photon imaging.

### Glioma induction in adult Camk2a-Cre-RiboTag or Thy1-EGFP-M mice

Isolated glioma cells were stereotactically injected in adult (10-12 weeks old) Thy1-EGFP-M (JAX ID 007788) or Camk2a-Cre-RiboTag mice. Homozygous RiboTag mice (JAX ID 011029) were crossed to Camk2a-Cre heterozygotes (JAX ID 005359) as described previously^39,92^. 5×10^4^ cells suspended in 1 μL OptiMEM (Gibco) were stereotactically injected to the right frontal lobe of adult mice using a Hamilton syringe with a 33 g needle, at a flow rate of 0.25 μL/minute (stereotaxic coordinates relative to bregma: 2.0mm anterior, 2.0mm lateral, 2 mm deep). Tumor growth was monitored by bioluminescent imaging as described previously^93,94^.

### Drug delivery

AZD8055 was dissolved in 30% (w/v) Captisol solution. A single dose (100 mg/kg) was administered to mice by oral gavage. Following six hours, mice were sacrificed, and tissue collected for molecular or histological analyses.

### Tissue processing for RNA isolation and Polysome Immunoprecipitation

Tissue was processed as described previously^39,92^. Briefly, the right frontal lobe of normal and glioma-bearing brains was flash frozen in liquid N2 before homogenization in 1 mL polysome lysis buffer (20 mM Tris-HCl pH 7.5, 250 mM NaCl, 15 mM MgCl2, 1 mM DTT, 0.5% Triton X-100, 0.024 U/mL TurboDNase, 0.48 u/mL RNasin, 0.1 mg/mL cycloheximide) at 4°C with a Dounce homogenizer. Homogenates were centrifuged at 14,000 x g for 10 minutes at 4° C. The supernatant was collected and half of it was used for total RNA extraction, performed using the RNeasy Mini kit (Qiagen), followed by ribosomal RNA depletion using Ribo-Zero rRNA Removal Kit (Illumina; MRZH11124) according to the manufacturer’s protocol. The remaining lysate was used for indirect IP of polysomes. First, SUPERase-In (0.24 U/mL) was added to prevent RNA degradation. Then, 15 μL of rabbit polyclonal anti-HA antibody (Abcam; ab9110) was added to the lysate and incubated at 4°C for 4 hours with rotation. 150uL of protein G-coated Dynabeads (30 mg/mL, Life Technologies) were washed 3x with 600 uL polysome buffer. The conjugated lysate was then added to protein G-coated Dynabeads and incubated with rotation at 4° C overnight. Beads were then washed 3x with 500 μL polysome buffer. RNA was eluted from the beads with polysome release buffer (20 mM Tris-HCl pH 7.3, 250 mM NaCl, 0.5 % Triton X-100, 50 mM EDTA) and extracted using the RNeasy Mini Kit. RNA integrity was assessed using a Bioanalyzer (Agilent).

### RNA sequencing libraries

Libraries were generated using the NEBNext Ultra Directional RNA Library Prep Kit for Illumina from New England Biolabs (E7420S) according to the manufacturer’s protocol. Nine RNAseq libraries were generated from each of the total RNA and polysome IP fractions. These included three tumor brains, three normal brains and three tumor brains from mice treated with AZD8055. Single-end sequencing of libraries was performed on a NextSeq 500 sequencer with a 75-cycle high-output kit (Illumina).

The RNA-Seq data were aligned to the mm10 genome and RefSeq transcriptome annotation using the STAR aligner^95^ and quantified using the featureCounts program in the subread package^96^.

### RNAseq data analysis

The neuronal enrichment score for each gene was calculated by performing DESeq2 analysis between the IP fraction and the total homogenate RNA from three independent tumor-bearing brains. Those genes with a positive Log2FoldChange and a padj<0.05 were considered to be enriched in IP fraction, representing transcripts associated with HA-tagged ribosomes from neurons. Differentially expressed genes between experimental conditions (control brain vs tumor-bearing brain and tumor brain vs AZD8055-treated tumor brain) were determined using DESeq2. Genes were further filtered based on their neuronal enrichment score to determine neuron-specific genes that were significantly up- or downregulated depending on the conditions.

### Gene Ontology Analysis

Gene ontology of the neuronally enriched and differentially expressed genes between control brain vs tumor-bearing brain and tumor brain vs AZD8055-treated tumor brain was performed using the PANTHER Classification System (https://www.pantherdb.org/).

### Gene Set Enrichment Analysis

All analyses were performed in pre-ranked mode using weighted statistics on the desktop GSEA software (version 4.3.2) downloaded from the Broad institute. Gene sets were compiled from Cahoy et al.^40^, Uezu et al.^48^, El Fatimy et al.^51^, Supplementary Table 1). In Figure 1a, a pre-ranked list of the significantly enriched neuronal genes derived from tumor brains was used. In Figures 1f and 1g, ranked lists from DESeq2 analyses comparing gene levels of genes that were both significantly enriched in the neurons and differentially expressed between tumor vs normal and tumor vs AZD8055-treated tumor were used. Ranking was based on the log2-fold change.

### Histological preparation of mouse brain sections and image analysis

At 5 weeks post-tumor cell injection or when luminescence reached a radiance of 10^6^-10^7^ (p/sec/cm^2^/sr), AZD8055 or vehicle were administered. After six hours, mice were anaesthetized using a mixture of ketamine and xylazine (100/10 mg/kg) and perfused with 10 mL PBS followed by 20 mL of 4% PFA solution at a rate of 5 mL/min. Brains were excised and post-fixed in 4% PFA for 24 hours. 40 μm - thick sections were obtained on a vibratome and were stained using a modified CUBIC-based protocol as detailed in Montgomery et al.^13^.

The following antibodies were used:

NeuN (Cell Signaling, #12943), HA (Millipore Sigma, #11867423001), mCherry (Abcam, #ab205402 and #ab167453), phospho-S6 (Ser240/244) (Cell Signaling, #5364), GFP (Thermo Fisher, #A10262), gephyrin (Synaptic Systems, #147008). All secondary Alexa Fluor conjugated antibodies were raised in goat and purchased from Thermo Fisher Scientific. All slices were stained with DAPI (Thermo Fisher, #D1306) and mounted on Superfrost Plus microscope slides (Fisher Scientific, #12-550-15) using Fluoro-Gel with TES buffer (Electron Microscopy Sciences, #17985-31).

Confocal images were acquired on a Zeiss LSM 800 confocal microscope using a Plan-Neofluar 40x/1.3 NA oil DIC objective. Fluorophores were excited using 405, 488, 561 and 639 nm lasers and captured on GaAsP detectors. Confocal stacks were acquired using 1 AU pinhole size. Images were exported to FIJI/ImageJ for further analysis.

For quantification of pS6 (Ser240/244) signal intensity in HA^+^ neurons, 4-5 independent fields from three biological replicates per condition were captured. Maximum intensity projection images were generated, and masks of the HA signal were created using the Otsu auto-threshold filter. Next, mean pS6 fluorescent intensity from each mask was calculated using the “Analyze Particles” function and results were displayed as mean fluorescent intensity per field.

For Gephyrin staining, an antigen retrieval step was performed using a citrate-based Antigen Unmasking Solution (Vector Laboratories, H-3300) by boiling in a microwave oven for 20 seconds. For analysis of gephyrin puncta, a minimum of 4 independent fields per condition from layer 5 of Thy1-EGFP-M mice were acquired. The gephyrin channel was auto-thresholded using the “Intermodes’’ method and particles were estimated using the “Analyze Particles’’ function for all masks with an area of 0.05-3 μm^2^ and circularity of 0.5-1.0. To identify the number of gephyrin puncta per EGFP^+^ neurons, masks of GFP were generated using the Shanbhag auto-threshold method. These were redirected to the gephyrin channel to identify particles using the above-mentioned threshold and size criteria.

For analysis of dendritic spine density in Thy1-EGFP-M mice, 200 μm-thick coronal sections were stained with antibodies against GFP and mCherry and mounted on glass coverslips. Confocal stacks of neurons and glioma cells were acquired at a frame size of 320×320 μm^2^ and a step size of 2 μm. Confocal images of secondary and tertiary dendrites were acquired at a frame size of 53.24×53.24 μm^2^ (1024×1024 pixels) and a step size of 0.4 μm. Confocal stacks were exported to Imaris (Oxford Instruments) for image analysis by an observer blinded to the conditions. Dendrites were traced manually, and an automatic intensity threshold was selected and processes with a maximum length of 4 μm and a minimum diameter of 0.3 μm for the spine head were chosen as size criteria defining dendritic spines. Before model generation and dendritic spine density calculation, a final manual step was performed by removal of erroneous signals deriving from adjacent filaments or inclusion of missed unambiguous dendritic spines deriving from the dendritic segment.

### DiOlistic labeling of brain sections and image analysis

DiI-coated tungsten particle preparation, delivery and tissue labeling were performed as described in Staffend and Meisel^97^. with minor modifications. Mice were anaesthetized using a mixture of ketamine and xylazine (100/10 mg/kg) and perfused with 10 mL PBS followed by 20 mL of a 2% PFA solution at a rate of 5 mL/min. Brains were removed, post-fixed for 10 minutes in 2% PFA and transferred to PBS. 200 μm-thick coronal sections were acquired on a vibratome (Leica). Sections were labeled by delivery of DiI-coated tungsten beads using a Helios gene gun (Biorad) at 120 psi. DiI was allowed to diffuse for 24 hours at 4°C followed by post-fixation with 4% PFA for 1 hour at room temperature. For antibody staining of DiI-labeled tissue, sections were permeabilized for 15 minutes in 0.1% Triton-X 100 in PBS and blocked in 0.01% Triton-X 100, 10% normal goat serum in PBS for 30 minutes at room temperature. Incubation of primary antibodies (anti-HA, HA11 mAb, Covance #MMS-101R; rabbit anti-Olig2, Millipore Sigma, #AB9610) was performed overnight at room temperature, followed by counterstain with secondary antibodies and DAPI for 1 hour at room temperature. Sections were mounted on glass slides and imaged on an inverted Zeiss LSM 800 confocal microscope using a 63x/1.4 NA or a 40x/1.3 NA oil immersion objective. Images of fields of labeled neurons were acquired at 1 μm steps. Z-stacks of secondary and tertiary dendrites were acquired at a frame size of 53.24×53.24 μm (1024×1024 pixels) and a step size of 0.2 μm. Dendritic spine number was counted manually by an observer blinded to the conditions with the help of FIJI software (NIH).

### APEX2 labeling and electron microscopy

A membrane-targeted construct of the pea ascorbate peroxidase (GAP43memAPEX2-FLAG) was PCR amplified from a pCAG vector and inserted in the first multiple cloning site of the pQCXIX retroviral plasmid at BamHI/EcoRI restriction sites. The resulting viral construct was used to infect p53^-/-^/PDGFA^+^/mCherry-Luciferase^+^ (APCL) cells, which were then injected in the subcortical white matter of adult (10-12 weeks old) Camk2a-Cre-RiboTag mice as detailed above. Once tumors reached appropriate size, mice were anesthetized and perfused for two minutes with carbonated normal Ringer’s solution (136 mM NaCl, 12.7 mM Na2HPO4, 50.3 mM KCl, 9.8 mM MgCl2, 59.5 mM NaHCO3, 20 mM CaCl2, supplemented with 0.2% w/v dextrose, 0.2 mg/ml xylocaine, 20 U/ml heparin) at a flow rate of 7 mL/minute. Mice were then perfused with fixative solution (4% paraformaldehyde, 0.5% glutaraldehyde, 2 mM CaCl2 in 150 mM cacodylate buffer pH 7.4) for 10 minutes. Brains were excised and post-fixed in fixative without glutaraldehyde for 24 hours before processing for serial sectioning scanning electron microscopy. serial block scanning electron microscopy (SBEM). Brain slices were prepared for SBEM as previously described^98^. To visualize APEX2-expressing glioma cells, the brain sections were exposed to diaminobenzidine (DAB (0.05%)/H2O2 (0.015%) in 0.1 M sodium cacodylate buffer until DAB labeling became visible under a microscope. Subsequently, the brain sections were washed with 0.1 M sodium cacodylate buffer. For SBEM osmium staining, the brain sections were incubated in OsO4 (2%)/potassium ferrocyanide (1.5%)/CaCl2 (2 mM) in 0.15 m sodium cacodylate for 1h. Then the brain sections were washed in water and transferred to filtered thiocarbohydrazide (0.5%) for 30 min. The brain sections were then washed in water and incubated in OsO4 (2%) for 30 min. The final steps involved dehydration using a series of ethanol and acetone solutions in six steps of 10 min each: 70% ethanol, 90% ethanol, 100% ethanol, 100% ethanol, 100% acetone, 100% acetone. Samples were then placed in a Durcupan ACM resin/acetone (1:1) solution overnight on a shaker. The following day, samples underwent two transfers into fresh 100% Durcupan ACM resin, six to seven-hours apart. After an overnight incubation in 100% resin, samples were embedded in fresh resin at 60 °C for a minimum of two days. The composition of Durcupan ACM resin (Sigma-Aldrich, St. Louis, MO) included 11.4 g component A, 10 g component B, 0.3 g component C, and 0.1 g component D. Acquisition and 3D reconstruction was performed as previously described^98^.

### Cranial Window Surgery and In Vivo Imaging

Cranial window surgeries were performed as previously described^13^, with some modifications. Briefly, adult Thy1-GCaMP6f mice (10-14 weeks) were anesthetized using isoflurane and injected with buprenorphine (0.1 mg kg^−1^ body weight), after which the dorsal skull was exposed and cleaned. A craniotomy was performed, and a single 4mm cranial window was placed with or without injecting ∼50,000 mCherry labeled glioma cells into the cortex (−1.5mm AP, 2mm ML, - 1mm DV). The glass window was glued in place using cyanoacrylate glue, after which a custom head plate was secured onto the skull using both cyanoacrylate glue and dental acrylic cement.

### In Vivo Two-Photon Imaging

To evaluate alterations in neuronal activity throughout tumor progression, we implanted tumor cells adjacent to the somatosensory cortex and performed longitudinal two-photon imaging of layer 2/3 excitatory neurons, which are known to respond to whisker stimulation^99^, in the whisker barrel cortex of awake Thy1-GCaMP6f mice. Mice were imaged awake and head-restrained in a custom-designed imaging enclosure at least 7 days following initial surgery. In the experiments where imaging was performed pre- and post-AZD8055, the pre-AZD8055 imaging was performed in the three hours immediately before drug administration and the post-AZD8055 imaging was performed in the three hours following oral gavage of 100 mg/kg AZD8055, starting approximately 30 minutes after drug administration. Imaging was performed under an upright ThorLabs Bergamo II two-photon microscope, running ThorImage LS and ThorSync, using a ×16 0.8 NA objective (Nikon) and a Ti-sapphire laser (Coherent) Vision-S) operating at 80 MHz with ∼85 fs pulses of light. The laser was set to 920 nm wavelength to image GcaMP6f fluorescence and to 1040 nm to excite mCherry fluorescence. Imaging was performed from the cortical surface to ∼250 µm deep at 30 Hz and 512 × 512 pixels covering 830 × 830 µm. For whisker stimulation trials, as described previously^66^ with modifications, a transparent acrylic rod was placed next to the right whisker pad at a fixed distance of 2 mm. Every stimulus trial consisted of a ten-second pre-stimulus epoch, a five-second whisker stimulation epoch, and a 15-second post-stimulus epoch. Six trials were grouped into a single run, separated by a 25-second intertrial interval, for a total of six whisker stimulation epochs in a single three-minute run. Whisker stimuli were applied by vibrating the bar at 25 Hz, a speed of more than 100 mm/s, and amplitude of more than 1 mm.

### Two-Photon Image Processing and Analysis

#### Field-Level Event Analysis

For each frame during a three-minute whisker stimulation runs, all the pixels in the imaging field-of-view (512×512 pixels) were averaged into a single mean value for each frame. This new average raw fluorescence time series used to calculate the ΔF/F and standard deviation of the three-minute run. To identify calcium peaks within each field-of-view, a minimum signal amplitude threshold *T* was employed, where *M* is the mean of all the non-zero values less than the 95th percentile of ΔF/F signal and *S* is the standard deviation of the values within *M*, and *T = M + 5*S. Single-Neuron Level Event Rate Analysis* Two-photon imaging data were first motion corrected and processed by Suite2P^62^. Regions of interest (ROIs) of neuronal soma were generated using Suite2P, using Cellpose add-on, and manually vetted for accuracy. ROIs and respective time courses (no neuropil subtraction) were exported to MATLAB, where further analysis was performed using custom scripts. We calculated ΔF/F for each pixel within each neuronal ROI using mean ROI intensity as the baseline. To identify calcium peaks within each neuronal ROI, a minimum signal amplitude threshold *T* was employed, where *M* is the mean of all the non-zero values less than the 95th percentile of ΔF/F data and *S* is the standard deviation of the values within *M*, and *T = M + 6*S.* For the neuronal event rate analysis in figures 3f and 4e, the entire three-minute whisker stimulation run was analyzed. For the neuronal event rate analysis in figure 4f, data was only considered from the five-second whisker stimulation epochs and the two-seconds immediately after the five-second whisker stimulation epoch. Any neuron with an event rate of 0 was excluded.

#### Single-Neuron Level Synchrony Analysis

To compute neuronal synchrony in a given imaging field-of-view, used in figures 3g and 4b, we used the methodology from Badimon et al.^63^ with slight modifications. Briefly, we took the neuronal events above threshold, as described above, and used the temporal location of these calcium events to identify each imaging frame as either containing an event or not. We then identified a “synchronous event” as any 500-millisecond interval where two or more different neurons in the same field-of-view had a calcium event. A 500-millisecond interval that contained only one neuron with more than one calcium event was not considered a synchronous event. For this analysis, only the five-second whisker stimulation epochs were considered. To correct for the variable number of neurons in each field-of-view, we reported the fraction of neurons involved in each synchronous event. To do this, the number of neurons demonstrating an event in a 500- millisecond interval were divided by the total neuronal ROIs in the field-of-view. Furthermore, since the analysis was focused on instances of elevated neuronal synchrony, the analysis was restricted to synchronous events two standard deviations above the mean of the control no-tumor data, which yielded ∼13% of neuron involvement. In other words, on average in control-no-tumor data during five-second whisker stimulation epochs, ∼5.4% of the neurons are recruited to a synchronous event with ∼3.8% standard deviation between whisker stimulation epochs. Therefore, the analysis was restricted to synchronous events demonstrating neuronal involvement two standard deviations above this typical control no-tumor level, yielding ∼13% neuronal involvement.

#### Single-Neuron Level Average Traces

To better estimate the temporal occurrence of each peak above baseline noise when averaging across individual neurons, used in figures 4g and 4h, each peak above threshold was temporally smoothed using a standard gaussian convolution function. To do this, the discrete event times of the events above threshold were convolved with a Gaussian kernel (300ms SD), thereby creating probabilistic event rates through time. Once each neuron’s events were smoothed by the gaussian convolution, the traces were summed across neurons within each millisecond to generate summed instantaneous probabilities of calcium event rates, and divided by the number of neurons, ultimately generating average event rates across a given number of neurons. Throughout the manuscript, these averages of gaussian convolved traces are referred to simply as “average calcium traces.” Modified from Merricks et al.^100^. In the modified raster plots and average traces (Fig 4g, 4h), the responses were included if there was a calcium event two seconds before stimulation, during the five-second stim, or two seconds after stim.

### Tumor Burden Calculation

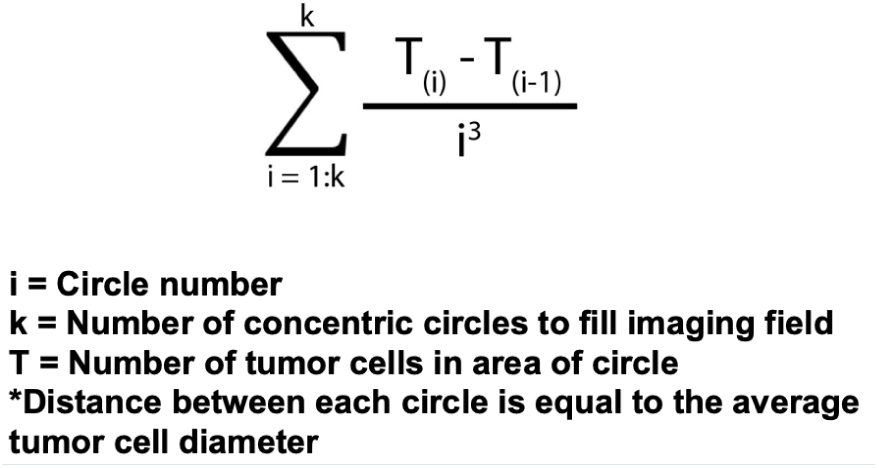

In order to calculate tumor-burden, image stacks of the mCherry tumor signal were taken for every field of view analyzed, beginning at the shallowest depth containing fluorescent tumor signal and ending at the deepest. Next, for each field of view, a maximum intensity projection across depth was calculated from these tumor image stacks and overlaid with the corresponding neuronal maximum intensity projection. ROIs for the tumor cells were generated using Cellpose^101^ and tumor-burden scores were calculated for each neuronal ROI using the equation below. For each neuronal ROI, equally spaced concentric circles were generated radiating outward from the ROI, and tumor cells were counted in each circle. Closer tumor cells were weighted more heavily in the computation, according to the equation below. Neurons were then split into low- and high-tumor-burdens based on the bimodal distribution of resulting tumor-burden scores.

### Cascade Detection

To detect cascades, we applied established methods from prior studies^58-61^. Within each frame, we converted the fluorescence signal of each pixel into binary form using a threshold of 5 times the signal’s standard deviation. In each frame, we identified clusters comprising at least 25 connected active pixels. This minimum threshold was set to ensure the detection of activities beyond that of a single neuron. Cascades were then defined as spatiotemporally contiguous clusters of active pixels^61^. A new cascade initiated when a cluster of previously inactive pixels became active and persisted as long as a connected cluster was detected in the next frame. Cascade size was determined by counting the total number of pixel activations during each cascade.

