## Supplemental_Figures for "Glioma-Induced Alterations in Excitatory Neurons are Reversed by mTOR Inhibition"

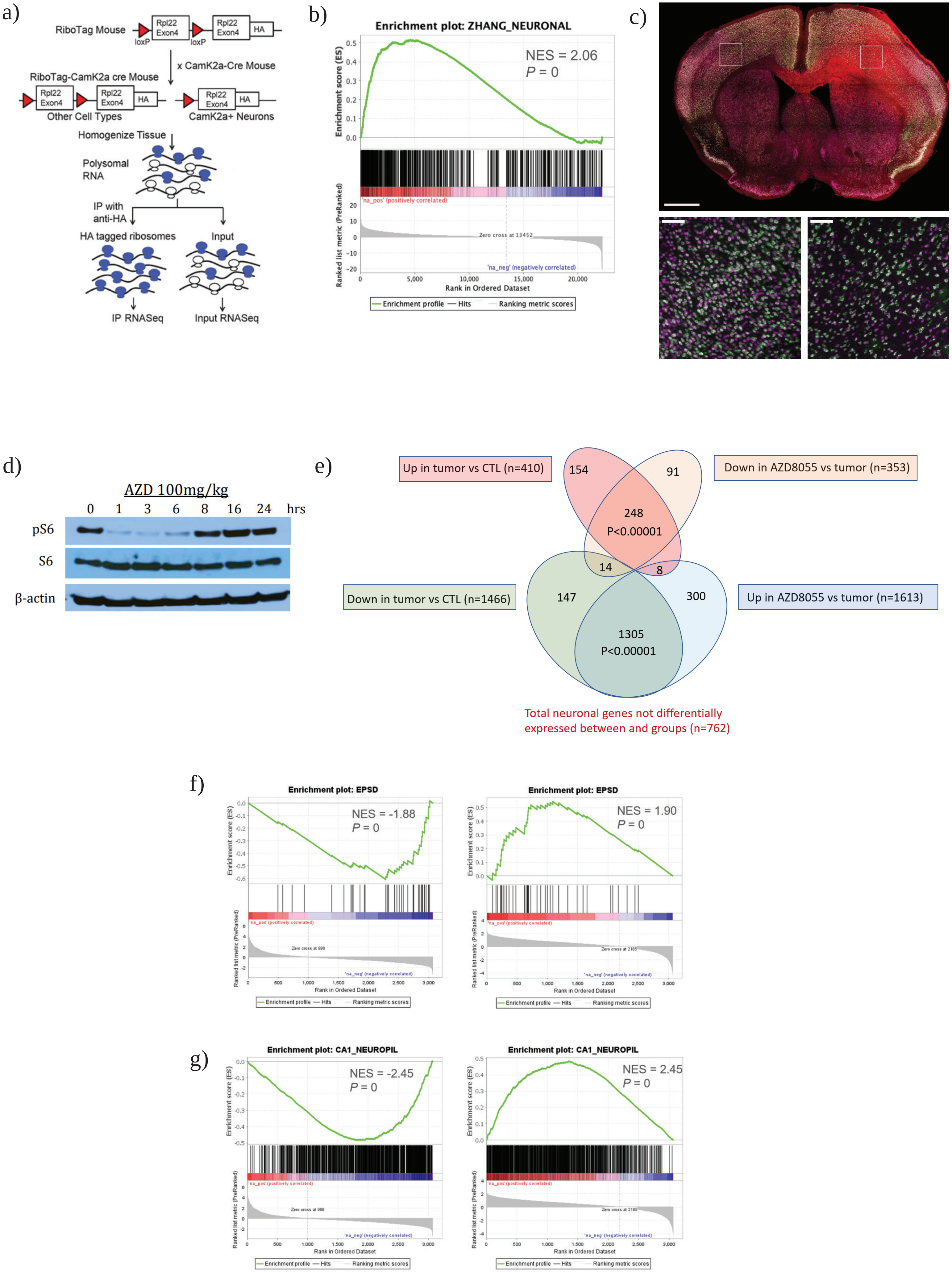
a)
RiboTag Mouse
loxP
loxP
Rpl22 Exon4
Rpl22 Exon4
HA
x CamK2a-Cre Mouse
RiboTag-CamK2a cre Mouse
Other Cell Types
CamK2a+ Neurons
Homogenize Tissue
Polysomal RNA
IP with anti-HA
HA tagged ribosomes
Input
IP RNASeq
Input RNASeq
b)
Enrichment plot: ZHANG_NEURONAL
NES = 2.06
P = 0
Enrichment score (ES)
Ranked list metric (PreRanked)
'na_pos' (positively correlated)
Zero cross at 13452
'na_neg' (negatively correlated)
Rank in Ordered Dataset
Enrichment profile
Hits
Ranking metric scores
c)
d)
AZD 100mg/kg
0 1 3 6 8 16 24 hrs
pS6
S6
β-actin
e)
Up in tumor vs CTL (n=410)
Down in AZD8055 vs tumor (n=353)
Down in tumor vs CTL (n=1466)
Up in AZD8055 vs tumor (n=1613)
154
91
248
P<0.00001
14
8
147
300
1305
P<0.00001
Total neuronal genes not differentially expressed between and groups (n=762)
f)
Enrichment plot: EPSD
NES = -1.88
P = 0
Enrichment plot: EPSD
NES = 1.90
P = 0
Zero cross at 998
Zero cross at 2185
Rank in Ordered Dataset
Enrichment profile
Hits
Ranking metric scores
g)
Enrichment plot: CA1_NEUROPIL
NES = -2.45
P = 0
Enrichment plot: CA1_NEUROPIL
NES = 2.45
P = 0
Zero cross at 998
Zero cross at 2185
Rank in Ordered Dataset
Enrichment profile
Hits
Ranking metric scores

**Supplemental Figure 1:**

a) Crossing of the RiboTag mouse to a Camk2-cre mouse results in RiboTag/Camkcre-2a mice in which Camkcre-2a+ neurons express HA tag in the ribosomal protein L22 (Rpl22HA). Following tissue homogenization, the polysomal fraction is either used for input for RNAseq or further immunoprecipitated using anti-HA antibody to enrich for the HA+ ribosomes (IP RNAseq).
b) GSEA plot showing enrichment of neuronal genes in the IP fraction of glioma-infiltrated brain. N = 3 mice per condition.
c) Decrease in the number of neurons in the ipsilateral hemisphere of glioma-bearing mice.
d) mTOR activity in response to AZD8055 in normal mouse brains. Non-tumor bearing mice were treated with 100 mg/kg AZD8055 for the time specified. Brain tissue from 0 hours was collected from untreated mice. N = 7 mice, 1 mouse per time point.
e) Venn diagram showing the significant overlap of neuronally enriched genes that are altered following AZD8055 treatment. N = 3 mice per condition.
f) GSEA demonstrates enrichment of excitatory synapse genes in tumor brain compared to control (left) and their depletion following AZD8055 treatment (right). N = 3 mice per condition.
g) GSEA demonstrates depletion of neuropil-expressed genes in tumor brain compared to control (left) and an enrichment following AZD8055 treatment (right). N = 3 mice per condition.

a)

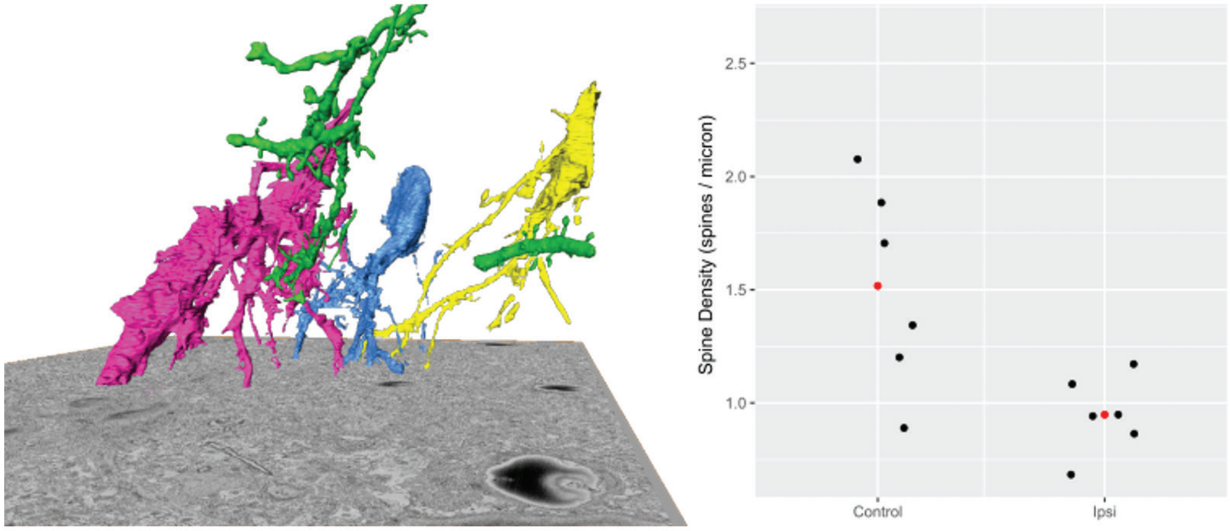
2.5
2.0
1.5
1.0
Spine Density (spines / micron)
Control
Ipsi

b)

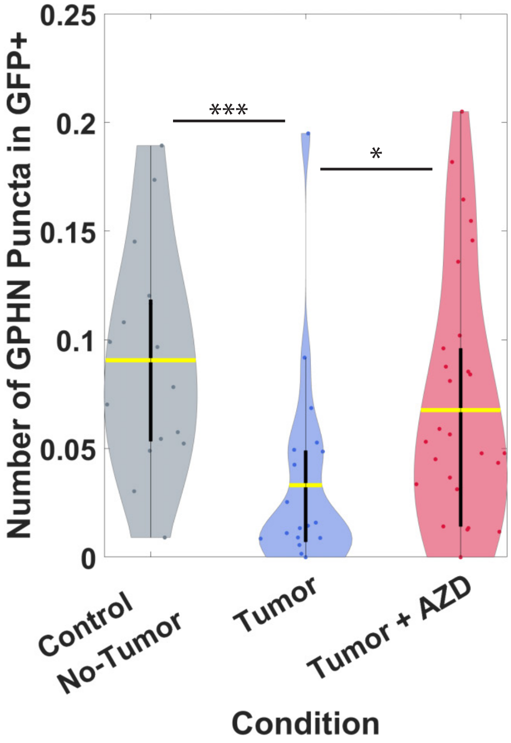
0.25
0.2
0.15
0.1
0.05
0
Number of GPHN Puncta in GFP+
***
*
Control
No-Tumor
Tumor
Tumor + AZD
Condition

**Supplemental Figure 2:**

a) Left; 3D reconstruction of dendritic segments (green) around APEX+ glioma cells (blue, yellow, and magenta) from serial scanning electron microscopy images. Right - plot of distribution of dendritic spines per µm of dendrite length from control brain and ipsilateral (glioma-infiltrated) cortex. Red dots represent the mean. N = 1 animal per condition.

b) Number of gephyrin puncta were quantified specifically localized inside Thy1-GFP+ neuronal ROIs. Gephyrin puncta were found to be significantly reduced in the tumor-associated GFP+ neurons compared to the no-tumor control, $p = 3.0118e-04$. In AZD8055 treated animals the gephyrin puncta inside GFP+ neurons were significantly higher than in the tumor alone, $p = 0.0232$. Wilcoxon rank-sum test was used. Yellow lines represent the mean. N = 18 imaging fields across 3 control mice, 29 imaging fields across 4 tumor mice, 30 imaging fields across 4 tumor + AZD8055 mice. Wilcoxon rank-sum test was used. Yellow lines represent the mean.

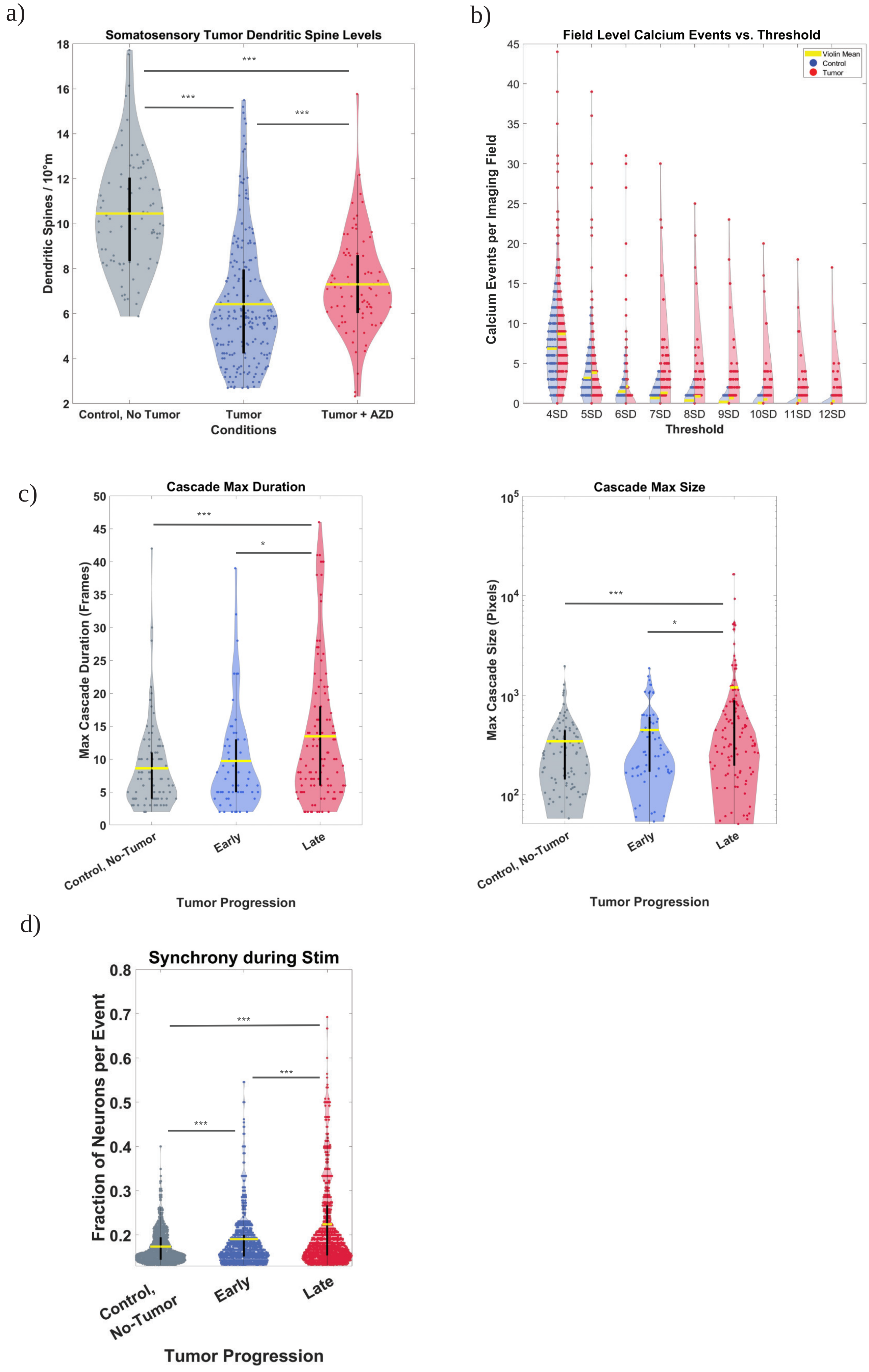
a)
Somatosensory Tumor Dendritic Spine Levels
Dendritic Spines / 10°m
Control, No Tumor
Tumor
Tumor + AZD
Conditions
***
b)
Field Level Calcium Events vs. Threshold
Violin Mean
Control
Tumor
Calcium Events per Imaging Field
4SD 5SD 6SD 7SD 8SD 9SD 10SD 11SD 12SD
Threshold
c)
Cascade Max Duration
Max Cascade Duration (Frames)
Control, No-Tumor
Early
Late
Tumor Progression
Cascade Max Size
Max Cascade Size (Pixels)
Control, No-Tumor
Early
Late
Tumor Progression
d)
Synchrony during Stim
Fraction of Neurons per Event
Control, No-Tumor
Early
Late
Tumor Progression

**Supplemental Figure 3:**

a) Glioma cells were implanted adjacent to the somatosensory cortex of Thy1-GFP+ mice, and dendritic spine densities were quantified across conditions. Dendritic spines were found to be significantly reduced in the tumor-associated GFP+ neurons compared to the no-tumor control, p = 6.0475e-22. In AZD8055 treated animals the dendritic sines were significantly higher than in the tumor alone, p = 9.1389e-05. The dendritic spine numbers of the AZD8055 treated tumor animals were still significantly lower than the control, 7.7266e-14. N = 79 dendrites across 3 control mice, 167 dendrites across 4 tumor mice, 79 dendrites across 4 tumor + AZD8055 mice. Wilcoxon rank-sum test was used. Yellow lines represent the mean.
b) For the field-level event analysis, first the mean ΔF/F in each field was taken and that single value was plotted for the entire three-minute run as a timeseries. Peak thresholds were determined based on standard deviations above the mean of the timeseries and the number of peaks above threshold in each run were computed. From this analysis we see that the mean number of peaks (horizontal yellow line) in the tumor is always higher than the control as the threshold increases in amplitude. This plot also illustrates that at certain thresholds above the mean (11, 12 standard deviations above the mean), there are no control events but there are still certain rare events in the tumor recordings. N=319 recordings across 8 tumor bearing mice and 124 recordings across 5 control mice.
c) Cascades were detected within each three-minute run of whisker stimulation, and the max duration and max size of the cascades within each three-minute run was plotted. The max cascade duration (frames) was significantly elevated in the late tumor recordings versus recordings from control no-tumor animals (p = 9.6146e-04) and between late recordings and early recordings (p = 0.0207). Similarly, the max cascade size (pixels) was significantly elevated in the late tumor recordings versus recordings from control no-tumor animals (p = 3.7465e-04) and between late recordings and early recordings (p = 0.0344). N=137 recordings across 7 early tumor mice, 182 recordings across 7 late tumor mice, and 124 recordings across 5 control mice. Wilcoxon rank-sum test was used. Yellow lines represent the mean.
d) The fraction of neurons involved per synchronous event was significantly higher in FOVs from later DPI recordings than from early DPI recordings (p = 3.2086e-25). The fraction of neurons recruited to a synchronous event was in early DPI recordings and later DPI recordings were both significantly higher than control recordings, early vs. CTL p = 3.1772e-13 and late vs. CTL p = 1.1836e-65. N = 2583 synchronous events across 6 early tumor mice, 3272 synchronous events across 7 late tumor mice, 2340 synchronous events across 5 control no-tumor mice. Wilcoxon rank-sum test was used. Yellow lines represent the mean.

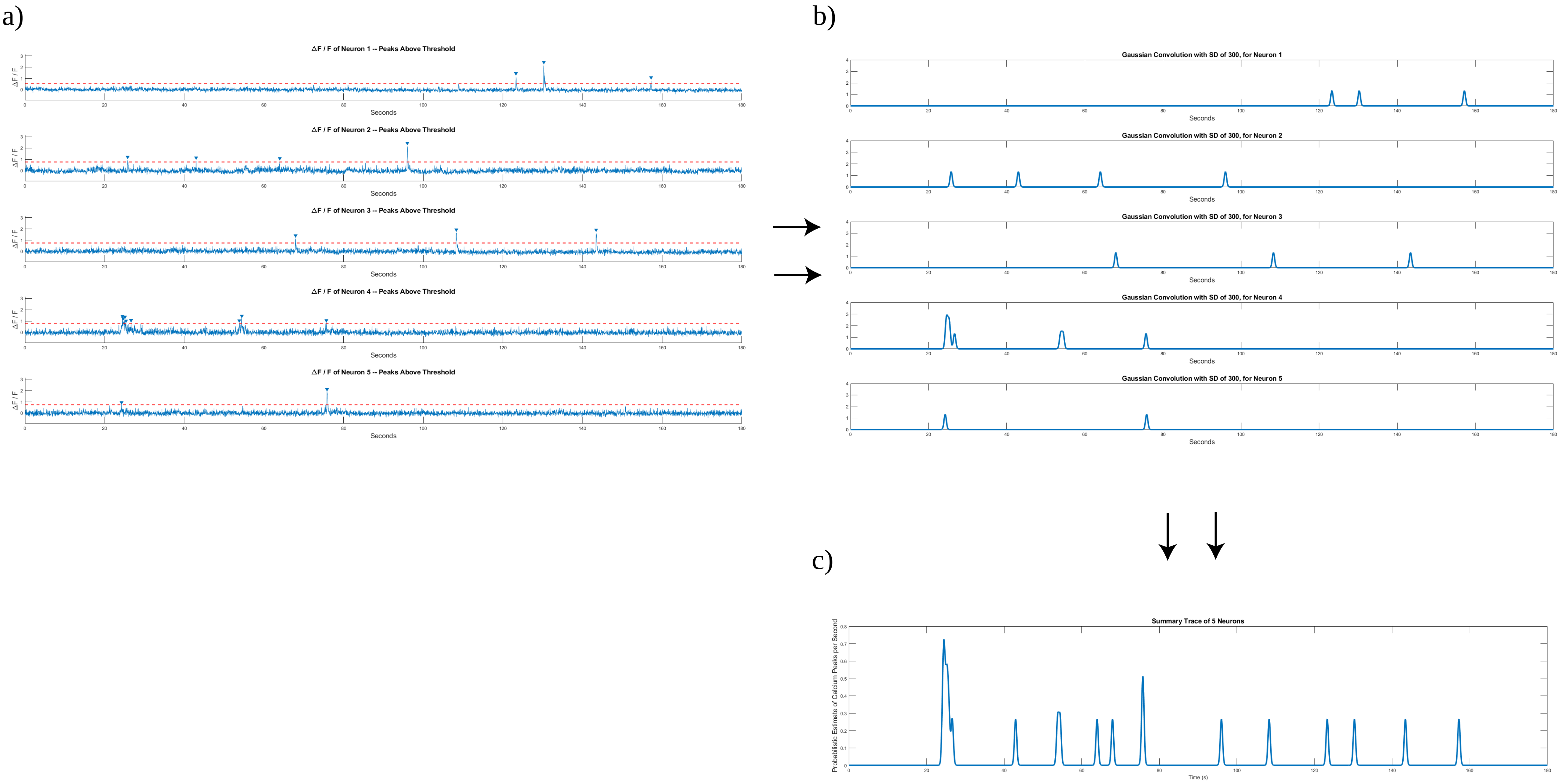
a)
ΔF / F of Neuron 1 -- Peaks Above Threshold
ΔF / F of Neuron 2 -- Peaks Above Threshold
ΔF / F of Neuron 3 -- Peaks Above Threshold
ΔF / F of Neuron 4 -- Peaks Above Threshold
ΔF / F of Neuron 5 -- Peaks Above Threshold
ΔF / F
Seconds
b)
Gaussian Convolution with SD of 300, for Neuron 1
Gaussian Convolution with SD of 300, for Neuron 2
Gaussian Convolution with SD of 300, for Neuron 3
Gaussian Convolution with SD of 300, for Neuron 4
Gaussian Convolution with SD of 300, for Neuron 5
Seconds
c)
Summary Trace of 5 Neurons
Probabilistic Estimate of Calcium Peaks per Second
Time (s)

**Supplemental Figure 4:**

a) Representative ΔF/F timeseries from 5 neurons, showing the events above threshold for each neuron.
b) Following temporal smoothing of peaks above threshold, using a gaussian kernel with a standard deviation of 300 milliseconds, each discrete event became a probability of a calcium event occurring in a period of time.
c) By summing the probabilities of calcium events across all neurons within each millisecond and dividing these summed instantaneous probabilities by the number of neurons, the average temporally smoothed event rate across all 5 neurons is computed.

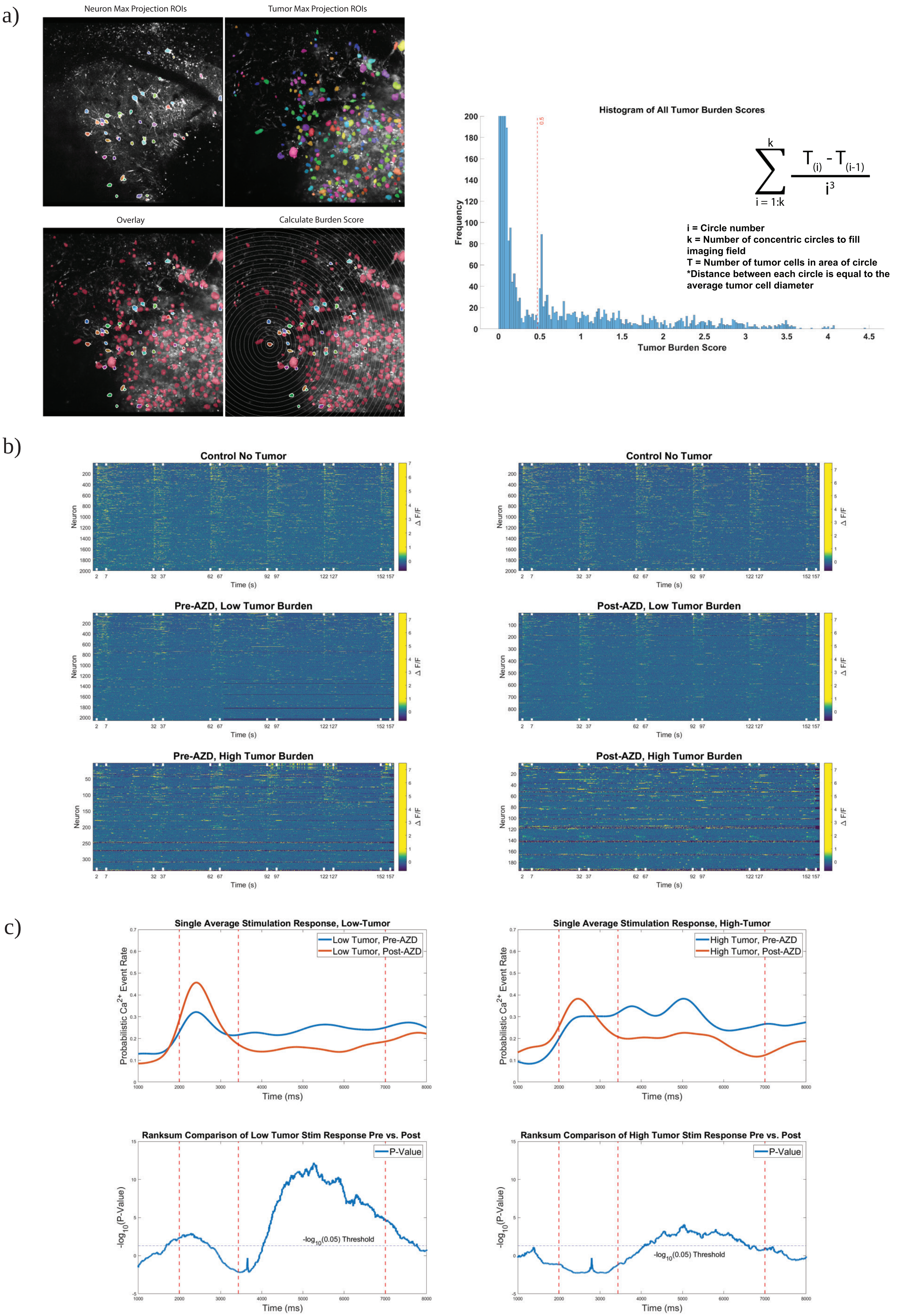
a)
Neuron Max Projection ROIs
Tumor Max Projection ROIs
Overlay
Calculate Burden Score
Histogram of All Tumor Burden Scores
Frequency
Tumor Burden Score
0.5
$\sum_{i=1:k}^{k} \frac{T_{(i)} - T_{(i-1)}}{i^3}$
i = Circle number
k = Number of concentric circles to fill imaging field
T = Number of tumor cells in area of circle
*Distance between each circle is equal to the average tumor cell diameter
b)
Control No Tumor
Pre-AZD, Low Tumor Burden
Post-AZD, Low Tumor Burden
Pre-AZD, High Tumor Burden
Post-AZD, High Tumor Burden
Neuron
Time (s)
Δ F/F
c)
Single Average Stimulation Response, Low-Tumor
Low Tumor, Pre-AZD
Low Tumor, Post-AZD
Single Average Stimulation Response, High-Tumor
High Tumor, Pre-AZD
High Tumor, Post-AZD
Probabilistic Ca2+ Event Rate
Time (ms)
Ranksum Comparison of Low Tumor Stim Response Pre vs. Post
Ranksum Comparison of High Tumor Stim Response Pre vs. Post
P-Value
-log10(P-Value)
-log10(0.05) Threshold

**Supplemental Figure 5:**

a) We employed Suite2P to generate neuronal masks and Cellpose to generate tumor cell masks based on the tumor cell mCherry signal. The maximum intensity projection across time was used for the neurons and the maximum intensity projection through the depth of the imaging field (Z-Stack) was used for the tumor cells. Then the tumor burden score was computed for each individual neuron. Based on the bimodal distribution of tumor burden scores, low tumor burden neurons were defined as neurons with tumor burden scores less than 0.5 and high tumor burden neurons had scores greater than 0.5. Included is the equation for such calculation for each neuron.

b) Modified raster plots were made by plotting color scaled ΔF/F of all the neurons from each respective neuronal population in order to visualize population activity. Data includes neurons from multiple tumor animals (n=5) and multiple controls (n=5), aligned according to the first whisker stimulation. The whisker stimulation start- and end-times are marked by the white notches along the horizontal axes. Neurons are sorted vertically according to the timing of their first event above threshold, with the earliest at the top. These plots demonstrate the clear temporal response of the control no-tumor neuronal populations to the stimulation onset, illustrated by the clear striping of high ΔF/F signal. This response is disturbed in the presence of both low and high tumor burden, before AZD8055 treatment, and by this visualization, appears to be qualitatively corrected in the low tumor burden group of neurons following AZD8055 administration.

c) Average stimulus evoked responses were computed for low tumor neurons (left plots) and high tumor neurons (right plots) before AZD8055 (blue) and after AZD8055 (red). Wilcoxon rank sums with Bonferroni-Holm correction for multiple comparisons were computed for each millisecond, comparing the likelihood of an event occurring in that millisecond between the two neuron groups. The horizontal line on the lower row of plots is the -log10(0.05) and the y-axis is the -log10(p-value), showing that the p-values plotted above that horizontal line are <0.05. The first and last vertical dashed lines indicate stimulus onset and offset. Evident from this analysis is that the main effect of AZD8055 is to reduce the neuronal activity in the later four seconds of the five-second stimulation epoch, this is best evidenced by the p-value reaching significance during the 2nd second of stimulation (~4000 milliseconds). N = 3741 pre-AZD8055 low tumor burden stim responses across 5 mice, 659 high tumor burden stim responses across 5 mice, 1611 post-AZD8055 low tumor burden stim responses across 5 mice, 342 post-AZD8055 high tumor burden stim responses across 4 mice.

**Supplemental Movie 1:** Field-Level Recording of Neuronal Activity at the Tumor Margin
Video of field-level events presented in Fig 3c, 3d, 3h. 30Hz imaging of neuronal calcium activity in Thy1-GCaMP mice during whisker stimulation, 134um below pial surface. Imaging field of view is 834x834µm (512x512 pixels) and Kalman filtered. Overlaid is the maximum intensity projection (Z-stack of pial surface to -200µm below, step size 2µm) of the mCherry+ tumor cells pseudo colored red.

**Supplemental Movie 2:** Single Neuron-Level Recording of Neuronal Activity at the Tumor Margin
Video of single neuron-level calcium activity. 30Hz imaging of neuronal activity in Thy1-GCaMP mice, 185µm below pial surface. Imaging field of view is 414x414µm (512x512 pixels) and Kalman filtered. Overlaid is the maximum intensity projection (Z-stack of -120 to -190µm below pial surface, step size 1µm) of the mCherry+ tumor cells pseudo colored red.
